# A VP2/3-derived peptide exhibits potent antiviral activity against BK and JC polyomaviruses by targeting a novel VP1 binding site

**DOI:** 10.1101/720318

**Authors:** Joshua R. Kane, Susan Fong, Jacob Shaul, Alexandra Frommlet, Andreas O. Frank, Mark Knapp, Dirksen E. Bussiere, Peter Kim, Elizabeth Ornelas, Carlos Cuellar, Johanna R. Abend, Charles A. Wartchow

## Abstract

In pursuit of effective therapeutics for human polyomaviruses, we identified a peptide derived from the BK polyomavirus (BKV) minor structural proteins VP2/3 that is a potent inhibitor of BKV infection with no observable cellular toxicity. The thirteen amino acid peptide binds to major structural protein VP1 in a new location within the pore with a low nanomolar *K*_D_. Alanine scanning of the peptide identified three key residues, substitution of each of which results in ∼1000-fold loss of affinity with a concomitant reduction in antiviral activity. NMR spectroscopy and an X-ray structurally-guided model demonstrate specific binding of the peptide to the pore of the VP1 pentamer that constitutes the BKV capsid. Cell-based assays with the peptide demonstrate nanomolar inhibition of BKV infection and suggest that the peptide likely blocks the viral entry pathway between endocytosis and escape from the host cell ER. The peptide motif is highly conserved among the polyomavirus clade, and homologous peptides exhibit similar binding properties for JC polyomavirus and inhibit infection with similar potency to BKV in a model cell line. Substitutions within VP1 or VP2/3 residues involved in VP1-peptide interaction negatively impact viral infectivity, potentially indicating the peptide-binding site within the VP1 pore is relevant for VP1-VP2/3 interactions. The inhibitory potential of the peptide-binding site first reported here may present a novel target for development of new anti-polyomavirus therapies. In summary, we present the first anti-polyomavirus inhibitor that acts via a novel mechanism of action by specifically targeting the pore of VP1.

## INTRODUCTION

BK polyomavirus (BKV), also known as human polyomavirus 1, is a small non-enveloped virus with a circular double-stranded DNA genome. BKV was first isolated from an immunosuppressed kidney transplant recipient in 1971 (Gardner et al., 1971), and is among the few clinically important human polyomaviruses, including JC polyomavirus (JCV) (Padgett et al., 1971) and Merkel cell polyomavirus (Feng et al., 2008). BKV is ubiquitous in human populations, with an estimated ∼80% sero-prevalence worldwide (Kean et al., 2009; Knowles, 2006). Primary exposure to BKV occurs in early childhood, with 50% of 3-year-olds and over 90% of 10-year-olds testing sero-positive (Knowles, 2001). Post-exposure, BKV infection is characterized by subclinical persistence with kidney tissue suspected as the viral reservoir (Ahsan and Shah, 2006; Heritage et al., 1981; Shinohara et al., 1993). Reactivation of BKV infection can occur in conditions of immunosuppression, particularly in the context of kidney and hematopoietic cell transplantation. BKV infection in kidney transplant recipients (KTRs) is first evident as viruria (20-70% of KTRs), which can progress to viremia (10-60%); BKV nephropathy (BKVN) is diagnosed in 3-4% of KTRs and 15-50% of those patients will suffer graft loss (Ambalathingal et al., 2017; Kuypers, 2012). The primary course of care for treating BKVN is reduction of immunosupressive therapy which carries the risk of acute graft rejection; up to 30% of BKVN cases treated by reduction of immunosuppressive therapy will experience an acute rejection episode (Bohl and Brennan, 2007; Sood et al., 2012). BKV reactivation in allogeneic hematopoietic cell transplant recipients can result in hemorrhagic cystitis (HC). In a recent study, 16.6% of allogeneic hematopoietic cell transplantations developed HC with BKV detected in the urine in 90% of cases (Lunde et al., 2015). There are currently no FDA-approved antiviral therapies for BKV, presenting an unmet medical need for these indications.

The lifecycle of BKV begins with virion binding to host GT1b and GD1b ganglioside (Low et al., 2006). The virus subsequently undergoes endocytosis via a caveolin-dependent pathway (Eash et al., 2004) and is trafficked in endosomes to the endoplasmic reticulum (ER) (Jiang et al., 2009; Moriyama and Sorokin, 2008), where a series of host cell enzymes orchestrate capsid disassembly (Goodwin et al., 2011; Schelhaas et al., 2007). The partially disassembled particle then interfaces with components of the ER-associated degradation (ERAD) pathway to undergo a critical step of retrotranslocation from the ER lumen into the host cell cytosol (Bennett et al., 2013; Jiang et al., 2009). Nuclear localization signal (NLS) domains within the capsid minor structural proteins then interact with components of the host nuclear pore complex to facilitate nuclear import of the viral genome (Bennett et al., 2015), wherein host cell transcription machinery initiates viral gene expression (Helle et al., 2017). The BKV virion is not known to contain any enzymes, viral or host (Fang et al., 2010); entry pathway steps are carried out via interactions between viral and host cell factors, and intra-virion interactions between the major and minor capsid proteins (Zhao and Imperiale, 2017).

The polyomavirus virion consists of a capsid formed by the major structural protein VP1 encapsidating the minor structural proteins VP2 and VP3, and the viral genome chromatinized with host histones (Cubitt, 2006). The capsid consists of 72 copies of homomeric VP1 pentamers cross-linked by intermolecular disulfide bonds to form a T=7d icosahedron structure (Nilsson et al., 2005). VP1 exists as a stable pentamer that contains a central pore, at the base of which a single copy of VP2 or VP3 is bound, forming a 5+1 complex as elucidated by X-ray and cryo-EM structures of infectious virions (Griffith et al., 1992; Hurdiss et al., 2018; Hurdiss et al., 2016; Liddington et al., 1991). All three structural proteins (VP1, VP2, and VP3) contain DNA binding domains (Clever et al., 1993; Soussi, 1986) and make contacts with the viral genome inside the infectious virion (Carbone et al., 2003; Hurdiss et al., 2016). VP2 and VP3 share a reading frame, with BKV VP3 consisting of the 232 carboxy-terminal residues of VP2 (Helle et al., 2017).

Reconstitution of the VP1 pentamer with full-length VP2 or VP3 has yet to be achieved; reconstitution of VP1 with a truncated VP2 protein and the corresponding X-ray structure has been reported for murine polyomavirus, implicating a “looping” structure for VP2 with the *C*-terminus interacting near the “base”, or inner virion-facing side, of the VP1 pentamer pore (Chen et al., 1998). Details of residues in the structural proteins contributing to the interactions of VP1 and VP2/3 have been elucidated primarily using either genetic (Bennett et al., 2015) or co-precipitation assays (Barouch and Harrison, 1994). Through these experiments, a region shared by both VP2 and VP3 near the carboxy-terminus of both proteins has been identified as required for the interaction with VP1 (Nakanishi et al., 2006). The biological function of the VP1 pore above the site of VP1-VP2/3 interactions at the base of capsid pentamers is unknown. For simplicity when referring to regions of the pore, “top” indicates the region nearest the exterior of the virus, and “bottom” or “lower” indicate the region nearest the interior of the virus.

In the current study, we report the discovery of a thirteen amino acid BKV VP2/3-derived peptide D1_min_ (corresponding to VP2 residues 290-302) that binds to BKV VP1 pentamers with single-digit nanomolar *K*_D_. We show that homologous peptide derived from JCV VP2/3 binds JCV VP1 with similar affinity, demonstrating a conserved binding interface. Protein-observed 2D NMR studies show this peptide interacts with VP1 residues in a previously uncharacterized location within the pore formed by pentameric VP1, with the binding location within the pore further corroborated by a structurally-guided model generated using X-ray data from co-complexed D1_min_ and VP1. Treatment of cells with D1_min_ in the context of BKV infection elicits nanomolar antiviral activity by the peptide, with a relationship established between peptide binding affinity to VP1 pentamers and antiviral potency. Additionally, we show the peptide exhibits antiviral activity against JCV, potentially indicating a pan-antiviral mechanism. We demonstrate through cell-based assays that the antiviral mechanism of action (MoA) involves blocking key steps in the viral entry pathway, likely prior to the critical step of ER-to-cytosol retrotranslocation. Mutations of residues in the VP1 pore that mediate peptide binding or of residues in the VP2/3 region from which the peptide is derived impact BKV infectivity, indicating the peptide-binding site may constitute a previously uncharacterized VP1-VP2/3 binding interface. In short, we report the first anti-BKV and anti-JCV molecule that directly targets the polyomavirus VP1 pentamer pore.

## RESULTS

### VP2/3-derived peptide binds pentameric capsid protein VP1 with high affinity

In order to better characterize the structural relationship between the BKV major structural protein VP1 and the minor capsid proteins VP2 and VP3, we focused on a stretch of amino acids near the carboxyl-terminus of VP2/3 previously referenced as “D1” (Nakanishi et al., 2006). The region is highly conserved among polyomaviruses, including JCV and simian virus 40 (SV40) (**Figure 1A; Supplemental Figure S1**). We initially tested binding of a 22-mer peptide VP2_281-302_ (APGGANQRTAPQWMLPLLLGLY; D1_22_) to purified BKV VP1 pentamers (VP1_2-362_) by Biacore surface plasmon resonance (SPR) and measured high affinity binding to the pentamer (*K*_D_ = 4.8 nM; **Figure 1B,C; Table 1**). The curve from a 1:1 interaction model overlays well with the SPR data (**Figure 1C**), consistent with a high quality, specific interaction, despite the hydrophobic nature of this peptide. In addition to directly measuring binding affinity by SPR, we developed an AlphaScreen assay to detect binding of carboxy-terminus biotinylated D1_22_ to VP1 and measure the half-maximal concentration at which the biotinylated peptide is displaced by unlabeled peptide (IC_50_) (**Figure 1D**, **Table 1**). The IC_50_ for unlabeled D1_22_ is 11±2.9 nM and this value is comparable to the SPR-determined *K*_D_. We additionally tested the homologous D1_22_ sequence from JCV with its cognate VP1 pentamer, and observed an IC_50_ of 44±6.4 nM (**Figure 1D**, **Supplemental Table S1**). Noting that protein-protein interactions often involve “molecular hot spots” where most of the binding energy is associated with a limited number of interactions (Van Roey et al., 2014), we split D1_22_ into two fragments, VP2_281-290_ (APGGANQRTA) and VP2_290-302_ (APQWMLPLLLGLY, henceforth referred to as D1_min_), and tested each fragment for binding to VP1. While no binding was observed for VP2_281-290_ up to 10 μM (data not shown), we observed similar binding affinity for the 13-mer peptide D1_min_ as was observed for D1_22_ (VP2_290-302_; *K*_D_ = 1.4±0.49 nM, IC_50_ = 3.6±0.57 nM) (**Figure 1D**, **Table 1**). Hereafter, references to amino acid positions in D1_min_ will be based on their sequence position in VP2.

**Table 1.**
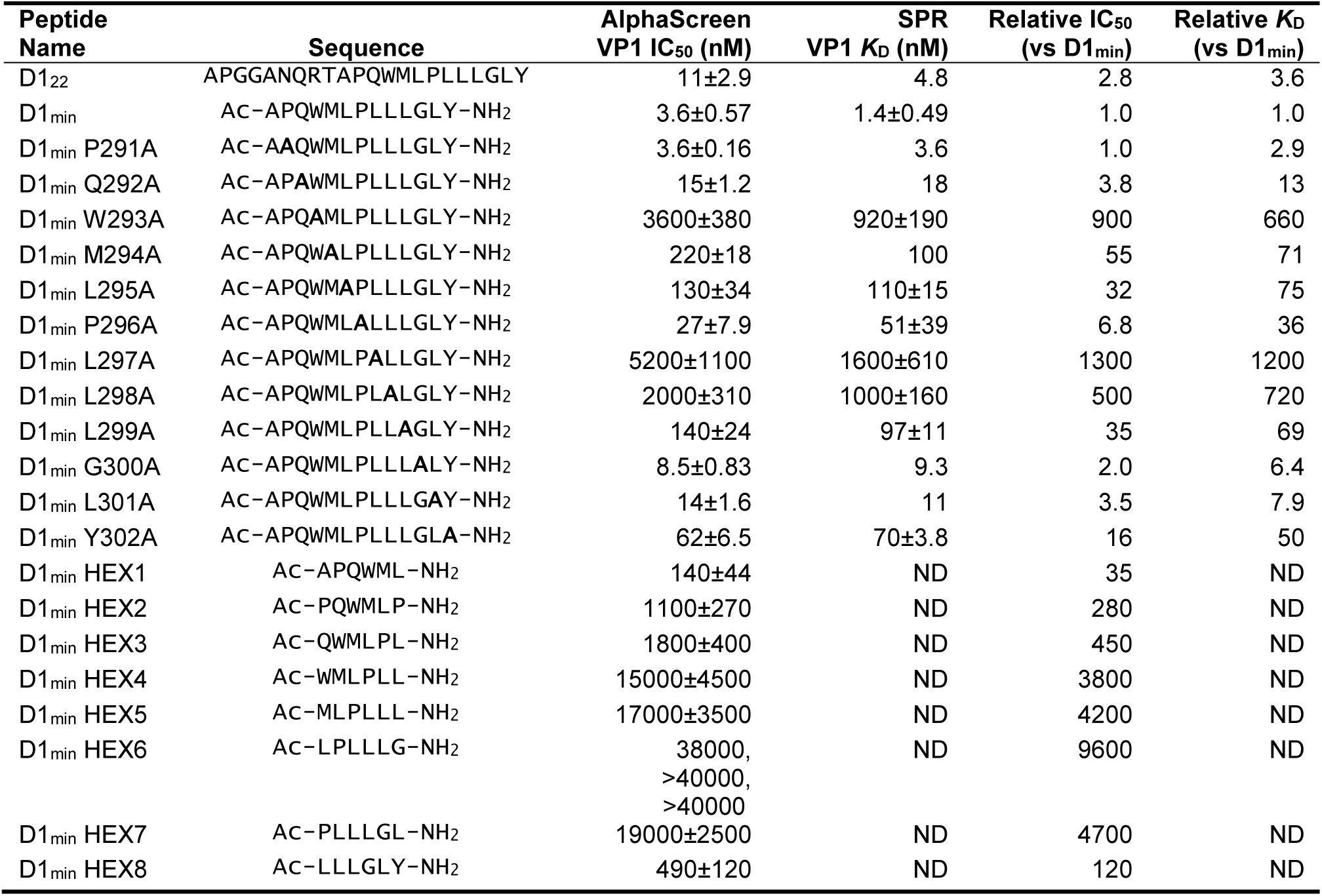
Peptide IC_50_ and *K*_D_ measurements. Values are mean ± SD where applicable (AlphaScreen: n=3; SPR: n=2). ND: Not determined.

**Figure 1.**
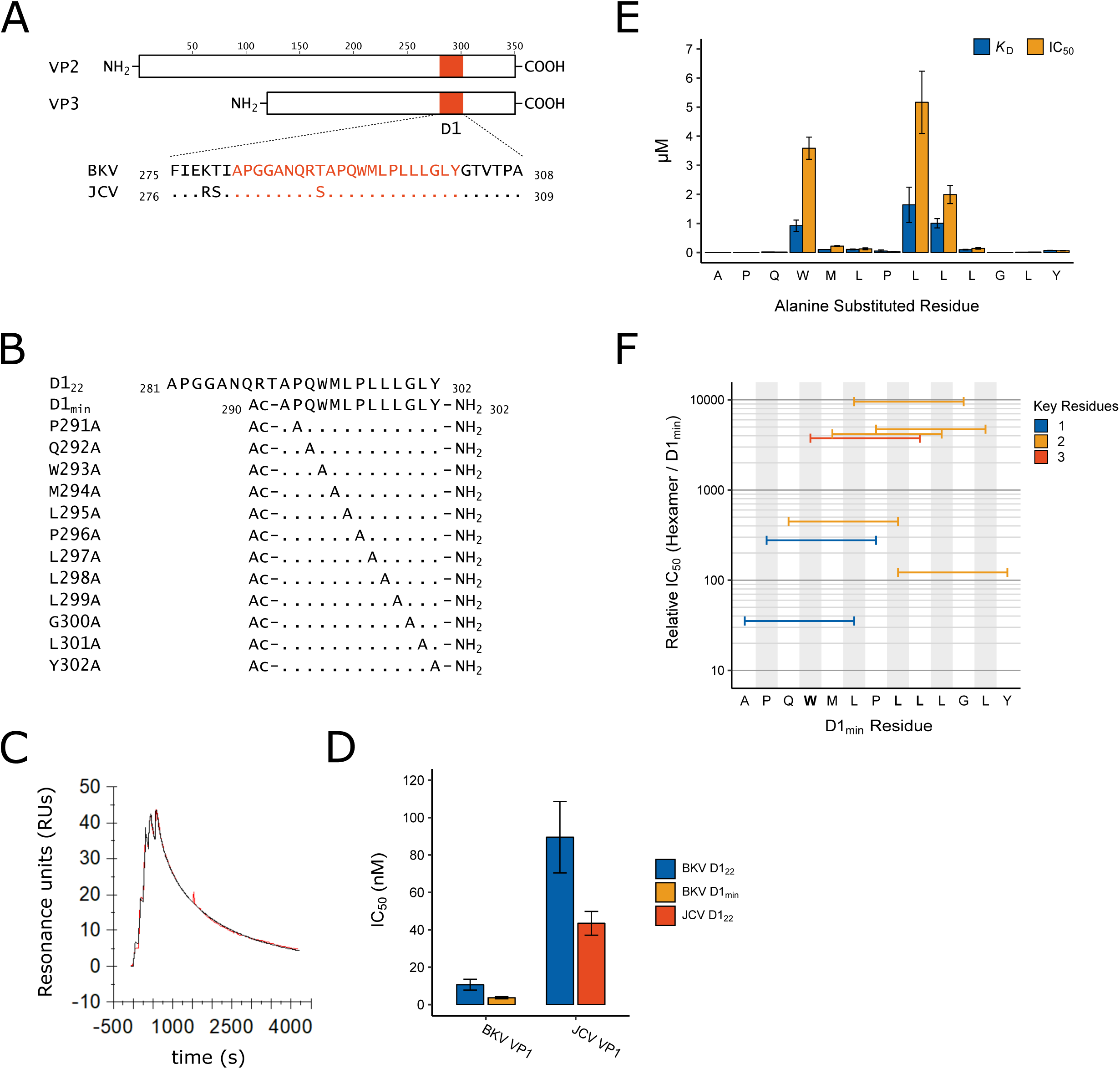
Identification of D1_min_ peptide and key residues contributing to interaction with VP1. **A.** Multiple sequence alignment of VP2/3 D1 region and flanking sequence. BKV: BK polyomavirus; JCV: JC polyomavirus. **B.** Sequence and index within BKV VP2 of peptides used in this study, highlighting alanine-scanning mutagenesis. Ac: acetyl group **C.** Representative surface plasmon resonance (SPR) sensorgram of single-cycle kinetic experiment showing association of D1_min_ with VP1 pentamer. Multiple (five) injections are shown, and dissociation of the peptide starts at peak response. Experimental data (red) and the 1:1 model of responses (black) are shown. **D.** Results of AlphaScreen competitive binding assay. Displacement of carboxy-terminal biotinylated D1_22_ peptide from either BKV or JCV VP1 was assayed using D1_22_ (BKV and JCV) or D1_min_ (BKV only), with IC_50_ concentration determined (mean ± SD, n=3 for BKV VP1, n=2 for JCV VP1). **E.** SPR-measured VP1 binding affinity (*K*_D_) and AlphaScreen displacement assay results (IC_50_; mean ± SD, n=3) for single-site alanine substitutions in D1_min_. **F.** AlphaScreen displacement assay IC_50_ values for D1_min_ rolling hexamer peptides (mean of n=3). Color indicates the number of key residues (W293, L297, or L298) present in the hexamer.

### Alanine substitutions in D1_min_ peptide reveal key residues contributing to D1_min_-VP1 interaction

To identify key side-chain residues involved in the interactions of D1_min_ with BKV VP1, we performed alanine scanning mutagenesis (Cunningham and Wells, 1989) on D1_min_, substituting one residue per peptide (**Figure 1B**), and analyzed the effect on binding to VP1 pentamers by Biacore SPR and the AlphaScreen competition assay (**Figure 1E**; **Table 1**). The SPR *K*_D_ and biochemical assay IC_50_ results are comparable for the alanine-substituted peptides (**Table 1**). Both assays identify residues W293, L297, and L298 as key determinants of high affinity binding, with each substitution causing ∼600-1000 fold loss of affinity to VP1 (*K*_D_ = 920±190 nM, 1600±610 nM, and 1000±160 nM, respectively). Alanine substitution of other residues (M294, L295, L299, Y302) results in 40-60 fold loss of affinity (*K*_D_), demonstrating that these may also contribute to binding affinity.

To see if we could further reduce the size of the peptide required for high affinity binding, we evaluated rolling hexamer peptides of D1_min_ in the AlphaScreen displacement assay (**Figure 1F**; **Table 1**). All hexamer peptides were significantly less potent relative to D1_min_. Notably, peptide D1_min_ HEX4 (_293_WMLPLL_298_) contains all three key determinant residues and has an IC_50_ that is greater than 1000-fold higher than that of full-length D1_min_ peptide. Rolling trimer peptides yielded similar results, showing greater than 1000-fold reductions in binding affinity to VP1 relative to the full-length D1_min_ peptide (**Supplemental Table S2**). We conclude that the key D1_min_ residues W293, L297, and L298 contribute significantly to the interaction of D1_min_ and VP1; however, additional peptide residues are required for the highest affinity binding.

### Peptide D1_min_ binds within the upper pore of VP1 pentamers

In order to determine the location of binding of the D1_min_ peptide to VP1 pentamers, protein-observed 2D-NMR spectroscopy was performed. ^1^H,^13^C-HMQC spectra of ^2^H,^12^C-BKV VP1_30-297_ with ^1^H,^13^C methyl-labeled residues Ile- (I), Leu- (L), Val- (V) and Thr- (T) were recorded in the absence and presence of increasing amounts of wild-type and W293A D1_min_ peptides, and ligand-induced chemical shift perturbations (CSPs) and line broadening were monitored. To enable mapping of binding locations we obtained peak assignments for select methyl groups through a combination of amino acid point mutations and a 4D-NOESY-HSQC based methyl walk (Proudfoot et al., 2016).

The NMR peaks assigned to BKV VP1 residues T224, T226, V231 (pro-*R* and pro-*S*), and V234 (pro-R and pro-S) were greatly affected by the addition of the peptides (**Figure 2A**). These residues are in close proximity to each other, clustering in the upper pore of the VP1 pentamer (**Figure 2B**). Binding of peptides to the upper pore appears specific since the site can be saturated (no additional CSPs observed at higher peptide concentrations) and only certain VP1 residues show CSPs while the majority of signals remained unaffected. As the set of perturbed peaks and the directions of chemical shifts are the same for both the wild-type and the alanine-substituted peptide, it can safely be concluded that both ligands have the same binding pose. Interestingly, while the wild-type peptide induces strong line broadening of certain peaks at sub-stoichiometric ligand concentrations, an observation that can be attributed to slow exchange kinetics, the alanine-substituted peptide causes pure chemical shift changes, which are usually a sign of fast chemical exchange (**Figure 2A**, top right corner). These results are consistent with our SPR and biochemical assay data, which showed that the wild-type D1_min_ peptide has a high affinity interaction with VP1 (low nanomolar *K*_D_) whereas the W293A peptide interacts with weaker affinity (*K*_D_ > 1 μM).

**Figure 2.**
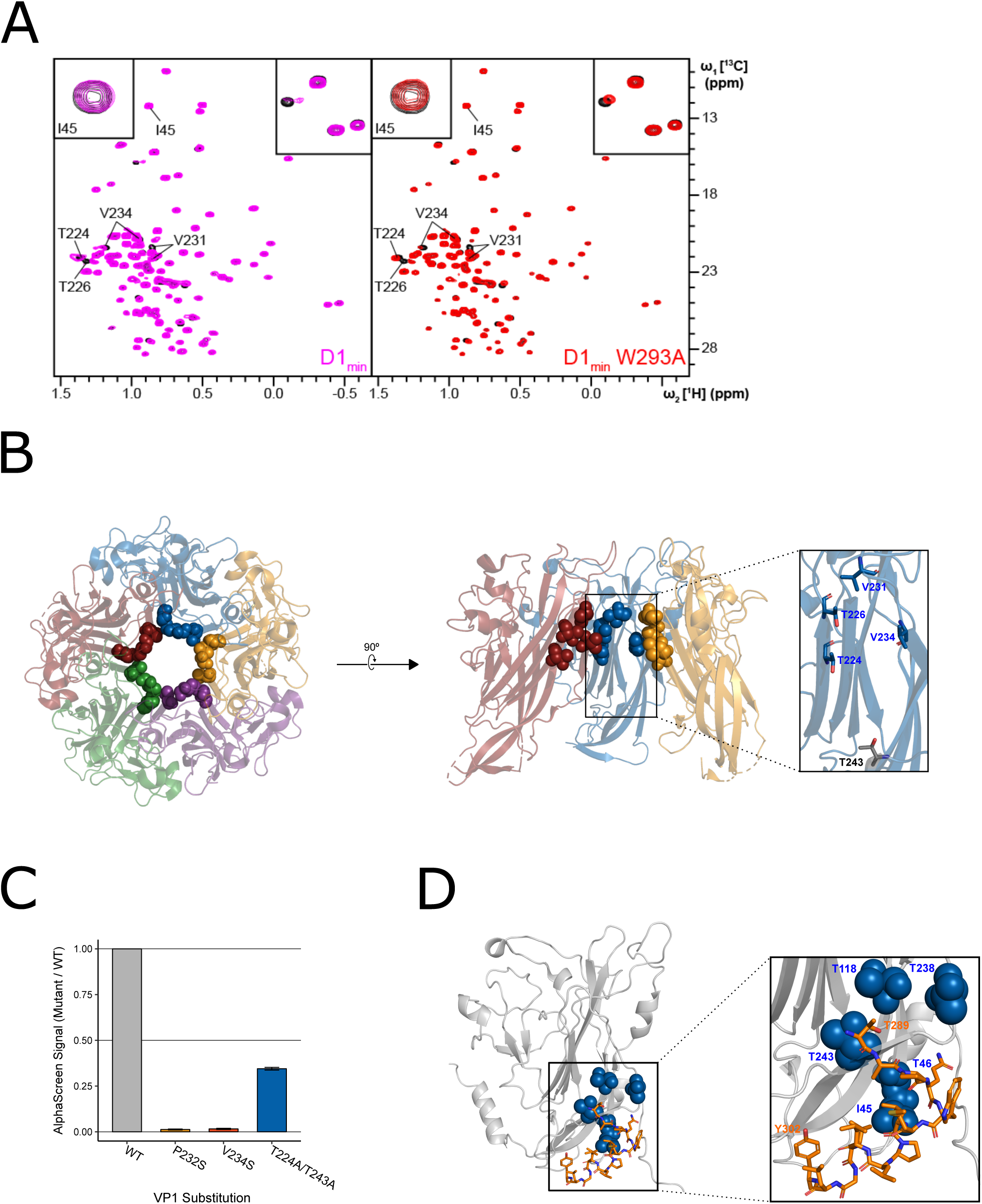
NMR characterization of the VP1-D1_min_ interaction. **A.** ^1^H,^13^C-HMQC spectra showing peptide (12.5 µM) induced perturbations of tr-VP1 (125 µM; black) ILVT methyl signals. **Left**: the wild-type D1_min_ peptide (magenta) causes CSPs and line broadening of peaks clustered in the upper pore of the target protein. The disappearance of peaks indicates slow exchange kinetics and thus, strong (usually sub-micromolar) binding (see inset in upper right corner). At sub-stoichiometric peptide concentrations no binding to a second site is observed as there are no changes of I45 (see inset in upper corner). **Right**: alanine-substituted W293A peptide (red) induces the same CSP pattern as the wild-type peptide, however, exchange kinetics are fast and no line broadening is observed. There is also no second site binding observed at low peptide concentrations. **B.** VP1 residues highlighted in **A** overlaid on X-ray structure of VP1 pentamer, looking down into the pore (left) and a cutaway side-view of three VP1 monomers (right). Spheres highlight VP1 residues that exhibit CSPs upon peptide binding (T224, T226, V231, and V234). Residue T243 is lower in the pore (shown in gray) and does not exhibit perturbations upon peptide binding (PDB: 4MJ1; Neu et al., 2013). **C.** Relative binding affinities of D1_22_ peptide to wild-type VP1 protein or VP1 containing pore residue substitutions using AlphaScreen detection method. Values are normalized to wild-type VP1 (mean ± SD). **D.** Overlay of “second-site” VP1 residues (I45, T46, T118, T238, T243; blue) on cryo-EM model of BKV VP1 (grey) and VP2 (orange) (adapted from PDB 6ESB, Hurdiss et al., 2018).

To validate the interaction of peptide D1_min_ within the VP1 pore, we focused on three sets of residues proximal to the observed CSPs, P232, V234, and T224/T243, and tested D1_min_ binding to VP1 proteins with substitutions at these residues using our AlphaScreen assay (**Figure C)**. We found that non-polar to polar substitution of either P232 or V234 lead to a substantial decrease in peptide binding relative to wild-type VP1 (P232S: 1.3±0.26%, V234S: 1.6±0.32% of wild-type signal). In contrast, a substitution that conserved the hydrophobicity of the putative binding site increased observed peptide binding (V234I: 250±26% of wild-type signal). Alanine substitution of VP1 pore residues further down into the pore (T224A/T243A) impact binding to a lesser degree (34±0.56% of wild-type signal). Consistent X-ray structures of VP1 proteins with these substitutions do not appear to have any major structural rearrangements (**Supplemental Figure S2A-B**), consistent with previous reports of polyomavirus VP1 pore mutants (Nelson et al., 2015), and we observe normal pentamer formation of these VP1 variants by size-exclusion chromatography during purification (data not shown). These results are consistent with a specific interaction of peptide D1_min_ within the upper pore of VP1 pentamers.

### NMR identifies second peptide binding site at the base of VP1

At a wild-type D1_min_ peptide concentration of 25 µM, well above the low nanomolar *K*_D_ observed for the primary peptide binding site in the upper pore, we observed additional ligand induced CSPs and line broadening in the 2D NMR spectrum. The VP1 peaks that were affected upon addition of peptide and for which assignments were available are I45, T46, T118, T238, and T243 (**Supplemental Figures S3A, S3B**). Without exception, these residues are located in the lower pore of the VP1 pentamer. Based on the first appearance of spectral changes at 25 µM D1_min_ peptide for a titration starting at 6.25 µM (where no CSPs were observed), we estimate that the *K*_D_ is greater than 250 µM. For peptide D1_min_ W293A, only at a ligand concentration of 100 µM did a few very weak additional peak shifts became visible; hence, the *K*_D_ value for this peptide is likely in the low single-digit millimolar range. At a wild-type peptide concentration of 50 µM and higher the above-mentioned peaks as well as signals from amino acids located in the upper pore show significant line broadening (**Supplemental Figure S3C**). This observation can potentially be explained by binding of multiple ligand copies or small soluble peptide aggregates. However, as the peptide induces signal perturbations of only certain VP1 residues all of which are in close proximity, the interaction site likely represents a binding hotspot. In line with this, the predicted location of the second interaction site is consistent with the modeled position of the D1 region of BKV VP2/3 in a recent cryo-EM structure (**Figure 2E**) (Hurdiss et al., 2018).

### X-ray model details interaction between D1_min_ peptide and VP1 pore

To further characterize the binding mode of D1_min_ within the VP1 pore, a structurally guided model was generated using 2.36Å resolution X-ray data from 13-mer D1_min_ peptide in complex with truncated VP1_30-297_ pentamers (**Supplemental Table S3**). The VP1 pentamer model is in good agreement with a previously published BKV VP1 pentamer structure (PDB: 4MJ1; Neu et al., 2013) (RMSD: 0.85Å; **Supplemental Figure S3D**). Electron density for the peptide is observed in the upper third of the VP1 pentamer pore, consistent with the NMR binding data (**Figure 3A**, **Supplemental Figure S3C**). Refinement with a best-fit model of observed electron density maps yields a primary chain of density consistent with an α-helical peptide running *N*-terminus at the top of the pentamer pore to *C*-terminus lower in the pore (**Figure 3B**), although electron density maps indicate multiple binding poses of the helix within the pore. VP1 pore residues that show peptide-induced CSPs by 2D NMR (T226, V231, V234; **Figure 2A-B**) as well as residues important for peptide binding as determined by substitution (P232, V234; **Figure 2C**) form a hydrophobic pocket around key D1_min_ residues L297 and L298 (**Figure 3C**). Interestingly, pocket structure appears to be largely unaltered by ligand binding (**Supplemental Figure S3D**). In conclusion, our structurally-guided model of D1_min_ in complex with VP1 agrees with NMR, alanine scan, and pore residue substitution studies placing the peptide in the upper pentamer pore and highlighting the importance of D1_min_ residues L297 and L298, as well as VP1 residues T226, V231, P232, and P234 in peptide binding.

**Figure 3.**
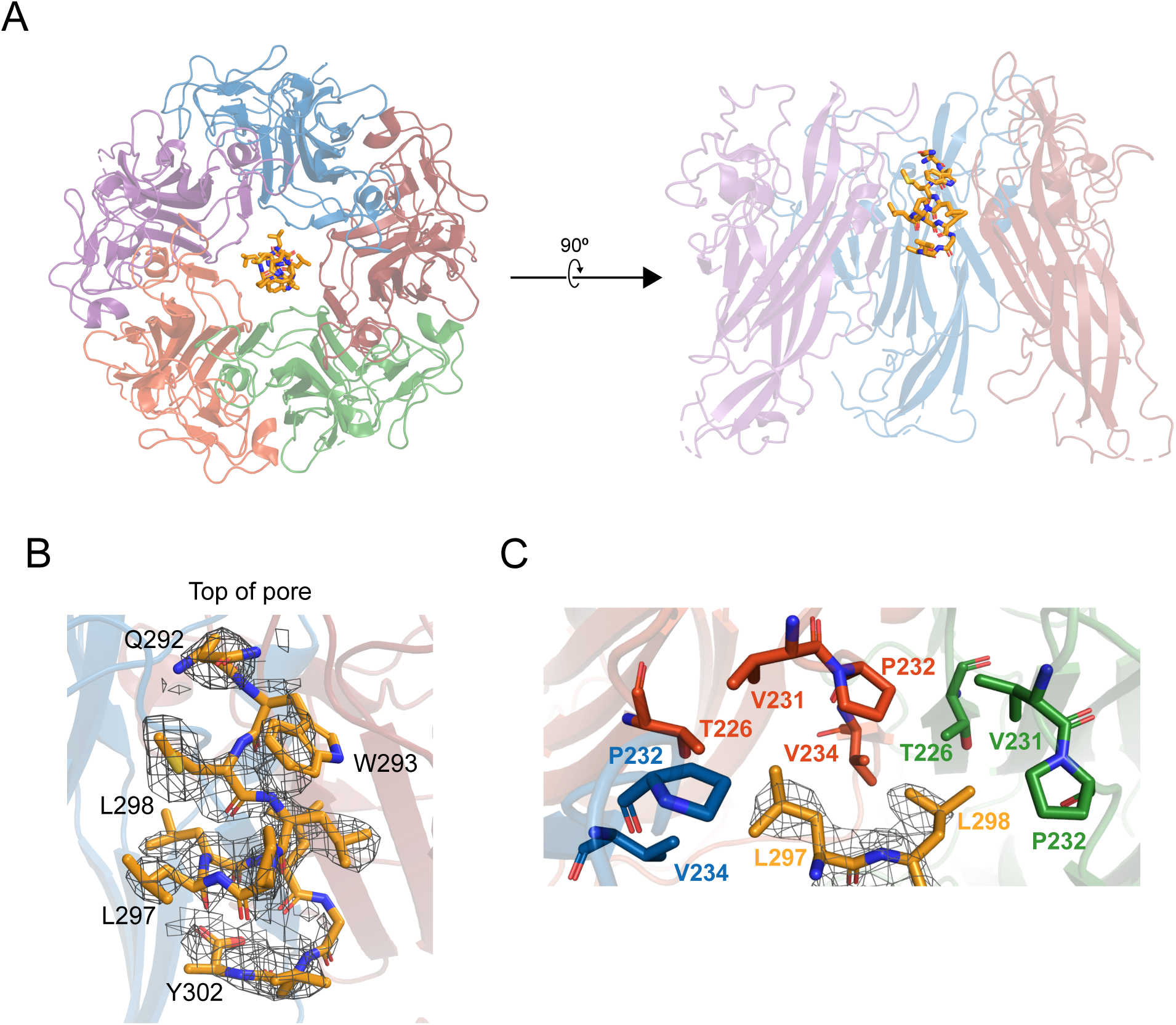
X-ray structurally-guided model of D1_min_-VP1 pentamer complex shows key residues mediating interaction. **A.** Structurally-guided model of structure of D1_min_ peptide bound to BKV VP1 pentamer. (Left) Top-down view of the model. (Right) Cutaway representation showing three VP1 molecules of the pentamer with D1_min_ peptide bound. **B.** D1_min_ 2Fo-Fc electron density map, contoured at 1σ, with model of peptide residues _292_QWLPLLLGLY_302_ built with guidance from the experimental maps. Start, end residues, as well as key binding residues W293, L297, and L298 are highlighted. **C.** Close-up of hydrophobic pocket formed by VP1 pore residues T226, V231, P232, and V234. Blue, orange, and green residues represent three distinct VP1 molecules within the pentamer. D1_min_ electron density for residues L297 and L298 (yellow), shown contoured to 1σ, correspond to regions of closest approach of the peptide to the pocket.

### D1_min_ peptide is a potent anti-BKV inhibitor

After observing high affinity binding of the D1_min_ peptide to the VP1 pentamer pore, we asked whether the peptide could inhibit BKV infection in a cell-based infectivity assay. Primary renal proximal tubule epithelial (RPTE) cells were pre-treated with a titration of peptide for 2 hours then challenged with infectious BKV (isolate MM), with indirect immunofluorescent staining for large T-Antigen (TAg) measured 48 hours post-infection (h.p.i) as a readout for productive infection. We observed potent antiviral activity from D1_min_ with a half-maximal effective concentration (EC_50_) of 30±6.6 nM without observable cytotoxicity in the concentration range tested (**Figure 4A,B**). Importantly, single alanine-substituted peptides of D1_min_ showed a loss of antiviral activity concordant with their loss of VP1 affinity in *in vitro* binding assays (W293A: >5000 nM, L297A: >5000 nM, Y302A: 280±51 nM).

**Figure 4.**
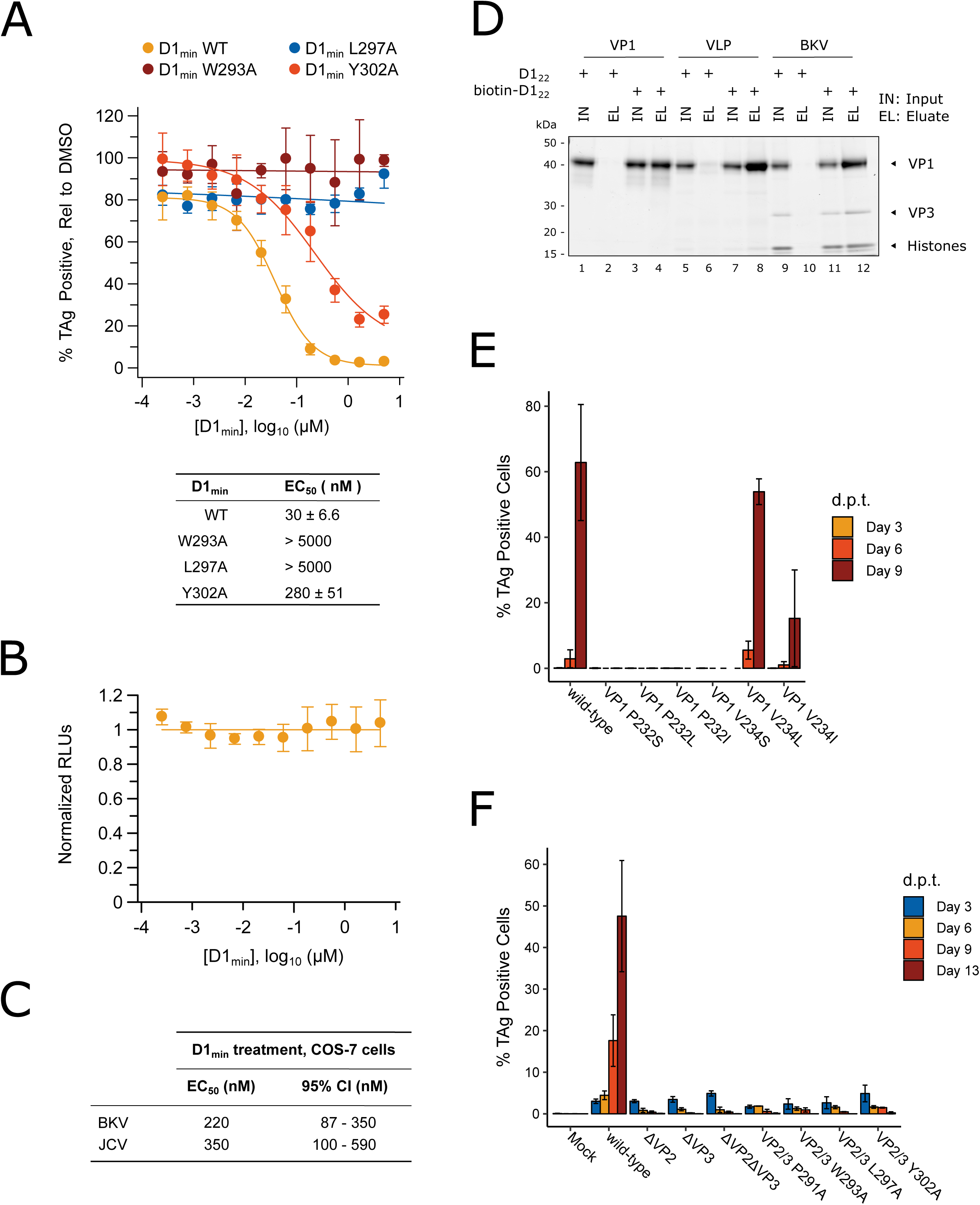
D1_min_ peptide has nanomolar antiviral activity. **A.** Dose-response curves for wild-type D1_min_ peptide and three alanine-substituted variants (W293A, L297A, L298A) in single-round BKV infection assay in RPTE cells (mean ± SD, n=3), and table of derived EC_50_ values. Productive infection is quantified by fraction of RPTE cells expressing BKV TAg by indirect immunofluorescent staining 48 hours post-infection (h.p.i). **B.** CellTiter-Glo luminescent cell viability assay to measure D1_min_ cytotoxicity in RPTE cells after two days of treatment. Relative light units (RLUs) are normalized to DMSO treatment (mean ± SD, n=2). **C.** D1_min_ EC_50_ values with 95% confidence intervals (CI) are shown for single-round infection assay of JCV and BKV in COS-7 cells, measuring fraction of VP1 expressing cells 72 h.p.i. **D.** Coomassie-stained gel showing streptavidin purification of VP1 pentamers, VP1 VLPs, or infectious BKV virions using either D1_22_ or biotinylated-D1_22_ peptide. **E.** BKV spreading infection assay with VP1 pore mutants, measuring TAg-positive cells 3, 6, and 9 days post-transfection of BKV genomic DNA. d.p.t.: days post-transfection (mean ± SD, n=2). **F.** Same as **E**, with BKV VP2/3 mutants (mean ± SD, n=3). While residue position is relative to VP2, VP2/3 indicates mutation is present in both proteins.

As the VP1 pore region and VP2 D1 region are highly conserved between polyomaviruses (**Supplemental Figure S1**), we further tested D1_min_ antiviral properties on the related human polyomavirus JCV. COS-7 cells were subjected to synchronized infection by either BKV or JCV (isolate MAD) followed by treatment with a titration of D1_min_ peptide, with indirect immunofluorescent staining for VP1 measured 72 h.p.i as a readout for productive infection (**Figure 4C**). We observe similar EC_50_ values for both polyomaviruses (BKV: 220 nM, JCV: 350 nM), albeit roughly 10-fold higher than observed for BKV in the RPTE cell model. This is likely due to the higher viral titers required for infection of COS-7 cells, as we have observed a positive relationship between BKV titers and measured D1_min_ EC_50_ (data not shown).

A notable difference between infectious BKV virions and VP1 pentamers or virus-like particles (VLPs) containing only VP1 is the presence of minor structural proteins at the base of the VP1 pentamer pore (Hurdiss et al., 2016). Based on the structural studies presented in **Figure 2,3**, the proposed mechanism of antiviral action by D1_min_ is through binding of the peptide to the VP1 pore. To confirm that D1_min_ peptide can bind to infectious BKV virions containing the minor structural proteins VP2 and VP3, we performed an affinity purification of biotinylated D1_22_ in the presence of VP1 moieties. Full-length VP1 pentamers, VLPs, or purified infectious BKV virions were incubated with 10-fold molar excess of either biotinylated or unlabeled D1_22_, followed by affinity-purification of biotinylated peptide and assaying co-purification of VP1. VP1 pentamers, VLPs, and infectious particles co-purified with D1_22_, demonstrating that the peptide can bind to infectious BKV virions (**Figure 4D**). Interestingly, only amino-terminal biotinylation was compatible with the assay; carboxy-terminal D1_22_ was unable to co-purify VP1 pentamers or VLPs, even when tested with truncated VP1 pentamers (VP1_30-297_) and extended peptides (**Supplemental Figure S4**, **Supplemental Table S2**). These data are consistent with the X-ray structurally-guided model placing the *N*-terminus of the peptide at the top of the VP1 pore. We conclude D1_min_ peptide can bind to infectious BKV virions that contain minor structural proteins at the base of VP1 pores.

### VP1 pore single-point mutants result in loss of BKV infectivity

As the D1 region of VP2/3 contains the same amino acid sequence as D1_min_, we tested whether residues that mediate D1_min_ binding to the VP1 pore are important for BKV infectivity. We performed site-directed mutagenesis of BKV *VP1* in the context of the viral genome, introducing substitutions at two key peptide binding residues in the VP1 pore, P232 and V234, and performed a spreading infection assay. Circularized wild-type or mutant BKV genomes were transfected into RPTE cells and productive, spreading infection was monitored by indirect immunofluorescent staining of expressed TAg over a time course of 3, 6, and 9 days post-transfection (d.p.t.) (**Figure 4E**). We observe robust spreading infection for wild-type BKV by 9 d.p.t. In contrast, BKV was completely intolerant of all tested substitutions at P232, as well as substitution V234S. V234L did not appear to affect BKV infectivity, and V234I, which showed increased binding to biotinylated peptide in an AlphaScreen biochemical assay, exhibited an intermediate phenotype with incomplete inhibition of viral spread. Importantly, all mutant viruses expressed similar levels of VP1 to wild-type BKV (**Supplementary Figure S2C**), dismissing interpretations that the observed phenotypes are due to differences in VP1 expression. Next, we performed reciprocal site-directed mutagenesis on BKV *VP2/3* in the context of the viral genome and repeated the spreading infection assay (**Figure 4F**). While wild-type and mutant BKV all expressed TAg at similar levels 3 d.p.t. after transfection, only wild-type BKV exhibited a spreading infection in culture. BKV was completely intolerant of VP2 or VP3 deletion, and of all tested alanine substitutions within the D1 region of VP2/3, with no detectable infectious virus produced from these mutant genomes. This is despite observing no significant impact on VP2/3 expression levels in mutants VP2 W293A and VP2 L297A (**Supplemental Figure S2D**). We conclude that residues involved in the VP1-D1_min_ interaction observed *in vitro* are required for productive BKV infection.

### D1_min_ peptide requires interaction with BKV for activity, but does not block viral endocytosis

Past studies have utilized broadly acting inhibitors of cellular activities to interrogate the polyomavirus entry pathway (Goodwin et al., 2011; Moriyama and Sorokin, 2008; Ravindran et al., 2017; Schelhaas et al., 2007). Such studies have been coupled with time-of-addition assays, in which treatment with inhibitors is initiated at different times during infection to correlate an inhibitor mechanism of action with a particular stage of BKV entry, including endocytosis (Eash et al., 2004), endosome maturation and vesicular trafficking (Eash and Atwood, 2005; Jiang et al., 2009), and ERAD / proteasome activity (Bennett et al., 2013). Similarly, we conducted a time-of-addition assay to better characterize at which stage of the BKV entry pathway D1_min_ antiviral activity occurs. RPTE cells were subjected to a synchronized infection at low multiplicity of infection (MOI) and inhibitor was added at varying times post-infection, with productive BKV infection assessed by indirect immunofluorescent staining of TAg expression at 48 h.p.i. (**Figure 5A**). In addition to treatment with D1_min_, we treated infected cells with an anti-BKV neutralizing monoclonal antibody P8D11 (Abend et al., 2017) and cell-penetrating TAT-tagged modifications (Vives et al., 1997) of D1_min_ which exhibit similar antiviral activity and biochemical potency to untagged D1_min_ peptide (**Supplemental Table S2,S4**). We observe a nearly complete loss of D1_min_ antiviral activity by 4 h.p.i. (**Figure 5B**), consistent with the timing of viral endocytosis (Eash et al., 2004). The BKV neutralizing antibody P8D11 parallels the time-dependent loss of activity of D1_min_. Cell-penetrating variants of D1_min_ show delayed loss of activity compared to the unmodified peptide, with only an approximate 50% loss of activity at 4 h.p.i. and a gradual tapering off of activity in subsequent timepoints. For comparison, previous time-of-addition work using Brefeldin A and nocodazole, treatments which affect viral trafficking to the ER, showed efficacy against BKV until 10-12 h.p.i (Jiang et al., 2009).

**Figure 5.**
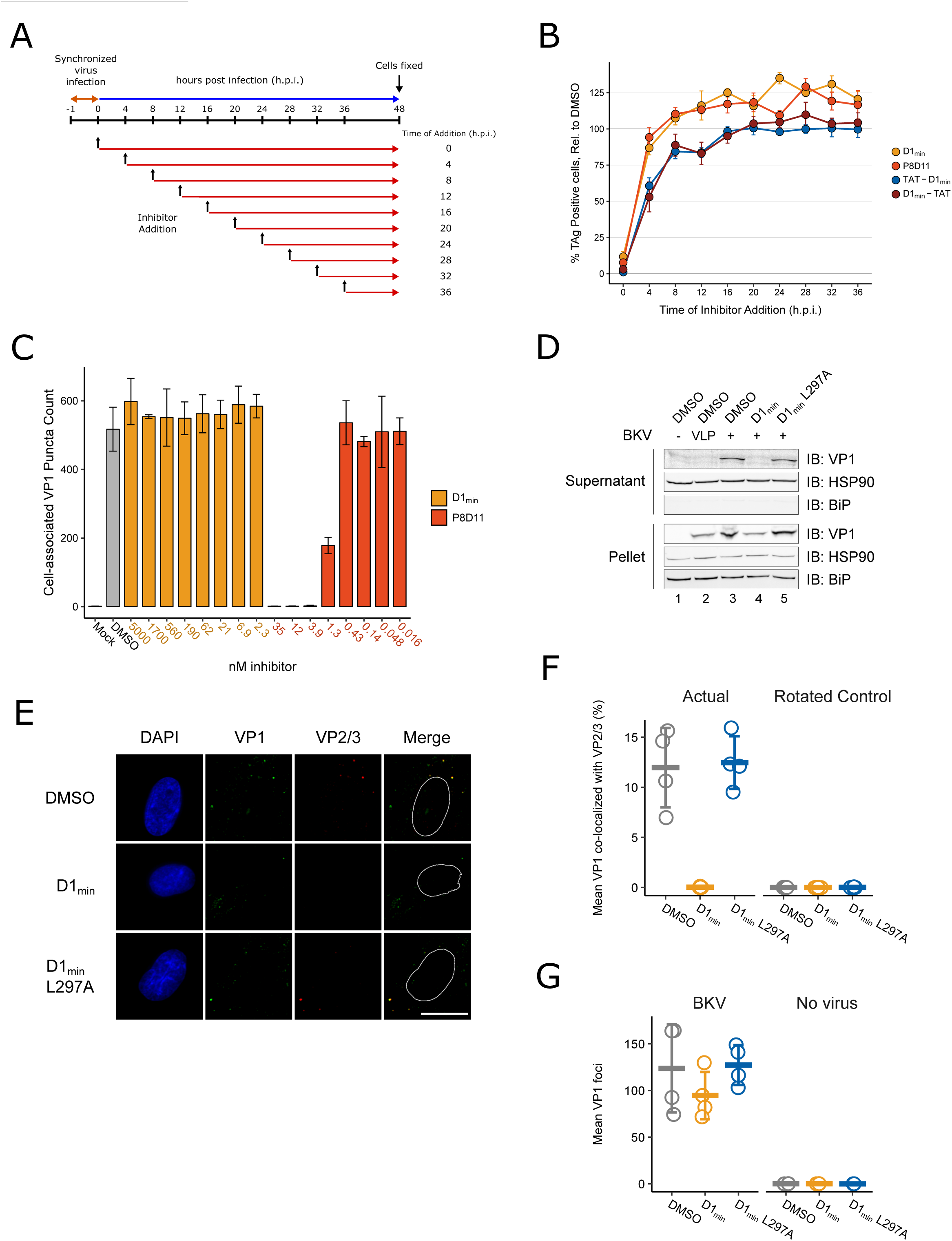
D1_min_ inhibits key steps in virion processing during entry **A.** Schematic of time-of-addition assay. **B.** Time of addition assay with D1_min_, cell-penetrating peptides TAT-D1_min_ and D1_min_-TAT, and anti-BKV neutralizing antibody P8D11. BKV infected cells were treated with inhibitors at 10-fold over measured EC_50_ concentrations. Productive infection is measured by the fraction of RPTE cells expressing BKV TAg by indirect immunofluorescent staining 48 hours post-infection (h.p.i), relative to DMSO-treated samples (mean ± SD, n=4). **C.** Virus cell binding inhibitor assay. BKV was treated with indicated inhibitor at indicated concentrations for 1 hour on ice, adsorbed to cells for 1 hour at 4°C, unbound virus washed away, and remaining cell-associated virus measured by indirect immunofluorescent staining of VP1 (mean ± SD, n=3). **D.** ER-to-cytosol retrotranslocation assay. RPTE cells subjected to a synchronized BKV infection (high MOI), cells were harvested 24 h.p.i, and lysates were fractionated into a supernatant (cytoplasmic) and pellet fraction. Fractions were then analyzed by SDS-PAGE, and VP1 protein and cellular compartment markers were detected by immunoblotting. **E.** Representative microscopy images of VP2/3 exposure assay. Minor capsid proteins were detected using a polyclonal antibody able to recognize both VP2 and VP3. Scale bar: 20 μm. **F.** Quantification of images exemplified in **E**, measuring fraction of VP1 stain co-localizing with VP2/3 stain, averaged per well. Rotated data indicates calculated co-localization between VP1 and VP2/3 stains after rotating VP2/3 images 90° to assess rate of random association between the two (mean ± SD, n=4). **G.** Quantification of VP1 foci in images exemplified in **E**, averaged per well (mean ± SD, n=4 for infected samples, n=2 for uninfected samples).

After observing loss of D1_min_ activity rapidly after initiating BKV infection in our time-of-addition study, we next asked whether D1_min_ acts as a cell-binding antagonist by performing a cell binding assay. Briefly, RPTE cells were incubated with BKV pre-treated with D1_min_ or P8D11 for 1 hour at 4°C to block endocytosis. Cells were rinsed, immediately fixed, and we performed indirect immunofluorescent staining for cell-associated VP1 puncta as a readout for cell-bound virions (**Figure 5C)**. We observed no effect on the number of cell-associated VP1 puncta in cells treated with D1_min_ up to 5 μM (>100-fold over observed EC_50_). In contrast, we observe loss of cell-associated VP1 puncta in the presence of the neutralizing antibody P8D11 treatment starting at concentrations >0.43 nM (>62 ng/mL), roughly the observed EC_50_ concentration. We conclude that D1_min_ does not block binding of BKV to cells and the observations from the time of addition experiment are due to the inability of the peptide to permeate the cell membrane rather than the peptide inhibiting BKV adsorption. This model is consistent with the delayed loss of antiviral activity observed for cell-penetrating variants of D1_min_.

### D1_min_ activity occurs prior to BKV ER-to-cytosol retrotranslocation

Next, we examined whether D1_min_ treatment affects the ER-to-cytosol retrotranslocation of BKV, a critical entry step and distinguishing feature of polyomaviruses (Dupzyk and Tsai, 2016). This transition can be assayed by fractionation of infected host cells and testing for the presence of VP1 protein in the cytosolic fraction (Bennett et al., 2013; Inoue and Tsai, 2011). RPTE cells were subjected to a synchronized BKV infection at high MOI followed by treatment of D1_min_ peptide (wild-type and L297A) at 10-fold over EC_50_ concentration. BKV VLPs were included as a negative control as they are unable to cross from the ER lumen into the cytosol (Geiger et al., 2011). At 24 h.p.i, cells were harvested, partially permeabilized with digitonin, fractionated between supernatant (cytosol) and insoluble pellet (e.g. ER, nucleus), and subjected to reducing SDS-PAGE followed by immunoblotting to detect the presence of VP1 in each fraction (**Figure 5D**). We observe the presence of VP1 in the pellet fraction for all samples treated with VLP or BKV, indicating the virus (or VLP) has undergone endocytosis under all treatments. In contrast, we only observe the presence of VP1 protein in the cytosolic fraction for untreated or D1_min_ L297A-treated BKV samples. Samples treated with wild-type D1_min_ or lacking the minor structural proteins VP2/3 (VLP) have no detectable VP1 protein in the cytosolic fraction, indicating the virus is unable to proceed through this step of the viral lifecycle.

### D1_min_ peptide inhibits exposure of minor structural proteins during capsid disassembly

During BKV entry, minor structural proteins which are initially concealed within the capsid become exposed, an event detectable by immunostaining (Norkin et al., 2002). Notably, inhibition of VP2/3 exposure is indicative of improper trafficking or disassembly of the virus (Bennett et al., 2013). We asked if D1_min_ affected this step of the BKV lifecycle. RPTE cells were subjected to a synchronized BKV infection at high MOI followed by treatment of D1_min_ (wild-type and L297A) at 10-fold over EC_50_ concentration. At 24 h.p.i., cells were fixed and stained by indirect immunofluorescence against VP1 and VP2/3 (**Figure 5E**). To assess the ratio of infectious particles to total particles in the cells, we calculated the fraction of virion particles (VP1-stained puncta) that co-localized with VP2/3 stain (**Figure 5F**). We observe a pronounced loss of VP1 co-localized with VP2/3 in wild-type D1_min_ treated cells as compared to our untreated control. In contrast, no change in co-localization between VP1 and VP2/3 is observed for treatment with the loss-of-function peptide D1_min_ L297A. VP1 staining was consistent across all samples (**Figure 5G**). We conclude that treatment with D1_min_ peptide results in virions that are unable to proceed through proper capsid disassembly that would result in the essential exposure of minor structural protein epitopes.

## DISCUSSION

Polyomaviruses are the causative agents of multiple human diseases, and the lack of effective antiviral therapeutics for the treatment of polyomavirus infections and associated diseases represent an unmet medical need. To identify potential therapeutics for BK and JC polyomaviruses, we explored the potential of targeting the major capsid protein VP1, one of the few proteins expressed by members of the polyomavirus family. This strategy of antiviral agents targeting viral capsids has also been explored for HIV, dengue virus, picornaviruses, and hepatitis B virus (Blair et al., 2010; Byrd et al., 2013; De Colibus et al., 2014; Deres et al., 2003; Fox et al., 1986; Klumpp et al., 2018; Lamorte et al., 2013).

We report the first described anti-polyomavirus inhibitor that acts through a novel anti-polyomavirus mechanism of action by binding the viral capsid pore. The BKV VP2/3-derived peptide D1_min_ binds specifically to major structural protein VP1 with high affinity (SPR *K*_D_ = 1.4 nM, biochemical IC_50_ = 3.6 nM) and NMR-based studies place the peptide binding site within the upper pore of VP1 pentamers. An X-ray structurally-guided model and residue substitution studies corroborate the peptide binding site within the VP1 upper pore. Interestingly, our NMR studies uncovered a potential second, weaker peptide-binding site at the base of VP1 pentamers, consistent with the modeled position of VP2/3 D1 region in previously published polyomavirus structures (Chen et al., 1998; Hurdiss et al., 2018). In cell-based assays, treatment with D1_min_ results in potent inhibition of BKV infection (EC_50_ = 30 nM) and elicits activity against the related human polyomavirus JCV. Using time of addition and fractionation assays to dissect the BKV entry pathway, our studies indicate D1_min_ mechanism of antiviral activity occurs sometime between endocytosis and retrotranslocation of the virus from the host cell ER into the cytoplasm. We can further refine the mechanism of action with the observation that D1_min_ treatment blocks exposure of minor structure proteins VP2/3 during capsid disassembly.

Our MoA studies utilized several strategies to help identify the stage of the viral lifecycle inpacted by D1_min_ treatment. We performed an inhibitor time of addition experiment that reported D1_min_ peptide acting very early in infection, concurrent with the timing of anti-BKV neutralizing antibody P8D11. However, in contrast to the neutralizing antibody, we clearly observed D1_min_ peptide treatment did not affect BKV binding to host cells, indicating a mechanism distinct from P8D11. Furthermore, cell-penetrating TAT-fused D1_min_ peptides continued to inhibit BKV infection at later addition time points as compared to the unmodified D1_min_ peptide, implying the observed rapid loss of activity of the unmodified D1_min_ peptide is likely due to cell impermeability rather than the timing of anti-BKV action. While observations for the TAT-tagged D1_min_ peptides also imply disruption of the early stages of BKV infection (e.g. entry and trafficking), there are caveats to direct interpretation of these cell-penetrating peptide data. First, the concentration of intracellular TAT-tagged D1_min_ peptide may be below the effective concentration required to inhibit infection when added at later time points post-infection, after the virus has been internalized. Second, the peptides may not localize to subcellular compartments where anti-BKV activity is required; previous studies on localization of TAT-tagged peptides and proteins demonstrate a predominant nuclear localization (Horton et al., 2008; Vives et al., 1997; Yang et al., 2002). Interestingly, not all small-molecule treatments that inhibit BKV entry result in the observed D1_min_ phenotype of blocking exposure of VP2/3 epitopes. Treatment with the proteasome inhibitor epoxomicin results in increased observed VP2/3 staining (Bennett et al., 2013), the opposite phenotype of D1_min_ treatment. Thus, blocking exposure of VP2/3 during viral entry may be specific to D1_min_ and not a general phenotype of anti-BKV treatments.

We characterized the binding determinants of D1_min_ as well as demonstrated binding specificity of the peptide to VP1 pentamers using alanine-scanning mutagenesis (Cunningham and Wells, 1989; Lozano et al., 2017). We identified three key residues in D1_min_ – W293, L297, and L298 – where single alanine substitutions resulted in ∼1000-fold loss of binding affinity to VP1 as measured in both an AlphaScreen competition assay and SPR studies. A structurally-guided model based on X-ray data acquired from VP1-D1_min_ complexes details an α-helical peptide binding in the upper VP1 pore, running *N*-to-*C* terminal from the top of the pore to the lower pore. Additionally, the model places key D1_min_ residues L297 and L298 within a hydrophobic pocket formed by VP1 pore residues T226, V231, P232, and V234. We observe these residues experience CSPs by 2D NMR upon peptide binding to VP1 (except the unlabeled P232). Biochemical peptide binding assays with substitutions of VP1 pore residues P232 and V234, as well as BKV spreading infection studies with mutations at VP1 P232 or V234, confirm and validate both the 2D NMR data and structurally-guided model of the peptide binding in the VP1 pore. Studies of trimer and hexamer truncations of the 13-mer D1_min_ peptide were unable to reconstitute high-affinity binding, including a hexamer peptide that contained the three key residues W293, L297, and L298, indicating these key residues are necessary but not sufficient for high affinity peptide binding to VP1. This may indicate that other residues likely contribute to peptide potency and/or the helical conformation of the peptide is important for its mode of binding. Peptides less than nine residues in length are unlikely to form secondary structures (Gellman, 1998; Manning et al., 1988). Accordingly, the high affinity binding of D1_min_ to VP1 likely uses some combination of structural and sequence elements.

As the D1_min_ peptide is derived from a native sequence found in the VP2/3 D1 region, and the peptide binds the VP1 pore with high affinity, intriguing questions are raised about the biology of the peptide and the pore to which it binds. Mutations in pore residues that are in close proximity to the peptide-binding site (VP1 P232, V234) generally result in noninfectious virus without grossly altering VP1 pentamer structure. Likewise, all tested mutations in the D1 region of BKV VP2/3 resulted in noninfectious virus. Both pore residues that we have shown by multiple methods are important for peptide binding (VP1 T226, V231, P232, V234) and the D1 region of VP2/3 from which the D1_min_ peptide is derived (VP2/3 290-302) are highly conserved across multiple polyomaviruses (**Supplemental Figure S1**). Viral proteins are subject to rapid sequence change over time due to high viral genome mutagenesis rates unless maintained by purifying selective pressure (Daugherty and Malik, 2012; Duffy et al., 2008; Kistler et al., 2007; Tokuriki et al., 2009); polyomaviruses are no exception (Buck et al., 2016; Pastrana et al., 2013). Thus, conservation of viral protein sequences is a strong indication of biological relevance. Further to this point, studies with JCV show both high affinity peptide binding to VP1 and inhibition of infection at a similar potency (EC_50_) as BKV in COS-7 cells, suggesting a conserved mechanism across at least two polyomaviruses. Interestingly, a study of JCV VP1 pore mutants found a deficiency in VP2/3 exposure (Nelson et al., 2015), corroborating our results with treatment of a pore-binding peptide. Both the conservation the residues in the VP1 pore and the VP2/3 D1 region that mediate D1_min_-VP1 interaction, and convergence of phenotype between the VP1 pore mutations and VP2/3 D1 region mutations, is suggestive that the peptide-binding site represents a previously uncharacterized VP2/3 binding site in the VP1 pore.

Multiple structures have been reported previously for infectious polyomavirus virions using both X-ray crystallography and cryo-electron microscopy (cryo-EM), with most reporting the minor structural proteins VP2/3 as a globular density at the base of the VP1 pentamer (Griffith et al., 1992; Hurdiss et al., 2016; Liddington et al., 1991). More recently, a cryo-EM structure of an infectious BKV virion mapped the resolved *C*-terminus of VP2/3, including the region containing the D1_min_ sequence, at the base of the VP1 pentamer (Hurdiss et al., 2018). These results are consistent with our current observations of a D1_min_ low affinity “second binding site” identified in the NMR studies.

An intriguing question is whether the high specificity and affinity of D1_min_ binding to VP1 is coincidental or reconstitutes an interaction between VP1 and the VP2/3 D1 region that occurs during the viral lifecycle. Dramatic restructuring of the viral capsid takes place during disassembly including reduction of intra-capsid disulfide bonds, decrease in virion size, and exposure of previously hidden minor structural protein epitopes from the capsid core (Bennett et al., 2013; Geiger et al., 2011; Inoue and Tsai, 2011; Jiang et al., 2009; Magnuson et al., 2005; Norkin et al., 2002). Our BKV infection assays indicate that residues involved in D1_min_ binding to the VP1 pore are critical for BKV infectivity. The low- and high-affinity D1_min_ binding sites found at the base and in the upper pore of VP1 pentamers, respectively, may represent VP1-VP2/3 interactions during different stages of disassembly, and subsequent exposure of VP2/3 may be required to allow for essential viral-host protein interactions. For example, interaction of the VP2/3 NLS with importin α/β is required for efficient nuclear import of SV40 and BKV viral genomes during entry (Bennett et al., 2015; Nakanishi et al., 1996; Nakanishi et al., 2002). The process by which polyomaviruses decrypt masked VP2/3 protein-protein interaction domains via structural rearrangements may involve the translocation of VP2/3 D1 region from a low-affinity binding site at the base of VP1 pentamers to the high-affinity binding site in the upper pore. Further studies are required to validate the proposed model, in particular studies that may deconvolute potential VP2/3 interactions with the lower and upper VP1 pore.

The essential step of membrane penetration by non-enveloped viruses is an incompletely characterized process (Kumar et al., 2018). Polyomaviruses complete this lifecycle step by exploiting the host cell ER-associated degradation (ERAD) pathway to undergo retrotranslocation from the ER lumen into the cytosol (Dupzyk and Tsai, 2016). This process involves viral interaction with numerous host factors and requires the presence of VP2/3 (Bagchi et al., 2016; Bagchi et al., 2015; Dupzyk et al., 2017; Geiger et al., 2011; Inoue and Tsai, 2017). Treatment of BKV with D1_min_ peptide inhibits two key observable lifecycle steps: exposure of VP2/3 epitopes to immunofluorescent staining and retrotranslocation of VP1 protein from the ER lumen into the cytosol. The MoA by which D1_min_ elicits theses antiviral phenotypes remains unclear. Of note is the observation that VP2/3 are required for retrotranslocation of virions out of the host cell ER lumen into the cytosol; VLPs (which lack minor structural proteins) are unable to complete this entry step (Geiger et al., 2011; Inoue and Tsai, 2011). We envision a few, non-exclusive models for D1_min_ antiviral action. D1_min_ binding in the VP1 pore may block interactions with host factors that bind there directly. Indeed, a cryo-EM study of infectious BKV particles identified possible heparin binding to the upper VP1 pore (Hurdiss et al., 2018). Blocking host factor interactions could result in improper trafficking of the incoming virion or inhibit capsid processing. Previous studies have shown that small molecules that inhibit polyomavirus trafficking to the ER block exposure of VP2/3 (Bennett et al., 2013; Bennett et al., 2015). In a second model, binding of D1_min_ peptide stabilizes the capsid and inhibits proper disassembly. The host restriction factor HD5 was shown to both inhibit VP2/3 exposure and stabilize the JCV capsid (Zins et al., 2014), suggesting that the two phenotypes may be related. Lastly, D1_min_ may disrupt an essential interaction between the VP1 pore and VP2/3 via the D1 region. Deciphering which of these models is correct will require further investigation using proximity-based methodologies to assess interactions with host factors and assays to monitor capsid disassembly.

In conclusion, we identified the first antiviral agent against BK and JC polyomaviruses that specifically targets the VP1 pore – the thirteen residue peptide D1_min_. The peptide is derived from polyomavirus minor structural proteins VP2/3 D1 region, and NMR and X-ray studies show the peptide binds to a novel site within the pore formed by pentameric VP1 capsid protein. The biological relevance of the interaction between peptide and the VP1 pore was confirmed by mutagenesis of the viral genome, with cell-based viral proliferation assays being impacted by mutations within the VP2/3 D1 region or the corresponding binding region within the VP1 pore. These observations indicate the peptide-binding site may be biologically-relevant, potentially constituting a previously uncharacterized VP1-VP2/3 binding interface. Given the single-digit nanomolar binding affinity of D1_min_ to the VP1 pore, the peptide provides a powerful new tool molecule for probing polyomavirus entry biology. Lastly, the inhibitory potential of the D1_min_-binding site within the VP1 pore first reported here may represent a novel target for development of first-in-class antiviral therapies to address the unmet medical need presented by polyomavirus infections.

## METHODS

### Cell culture

Primary renal proximal tubule epithelial (RPTE) cells were purchased from ATCC (PCS-400-010) and cultured in RenaLife Basal Medium with supplements (Lifeline Cell Technology LL-0025) as previously described (Abend et al., 2007). COS-7 cells were purchased from ATCC (CRL-1651) and cultured in DMEM medium (Corning Cellgro 10-017-CV) supplemented with 5% fetal bovine serum (FBS) (Seradigm). HEK-293 cells were purchased from ATCC (CRL-1573) and cultured in DMEM medium (Corning Cellgro 10-017-CV) supplemented with 10% FBS (Seradigm). Cells were cultured at 37°C with 5% CO_2_.

### Virus stock generation

BKV stocks were generated by transfection and infection of cells as previously described (Abend et al., 2007). Briefly, BKV ST1 MM viral genome was excised from pBR322 plasmid (ATCC 45026) using *BamHI* (NEB), cleaned up using QIAquick PCR Purification Kit (Qiagen), and re-circularized using T4 DNA ligase (NEB) overnight at 16°C. The re-ligated viral genomes were extracted using phenol:chloroform:isoamyl alcohol (25:24:1,v/v) (Sigma) and aqueous phase was separated using Phase Lock Gel Heavy tube (5Prime), followed by ethanol precipitation and resuspension of viral genomes in Buffer EB (10 mM Tris-HCl, pH 8.5, Qiagen). HEK-293 cells were transfected with 2-4 μg of viral genome using Lipofectamine-2000 (Invitrogen) and Opti-MEM (Gibco) according to manufacturer’s protocol, and cells were cultured for 10-14 days until CPE were observed. Cells were freeze-thawed three times and supernatant cleared by centrifugation at 1600 rpm for 15 minutes. Low-titer virus from resulting supernatant was used to infect either RPTE or HEK-293, and cells were cultured for 12-14 days (RPTE cells) or 21-28 days (HEK-293 cells) until CPE was observed. Cells were then scrapped, freeze-thawed three times, and purified as described below.

JCV stocks were prepared similarly. The genome of JCV genotype Ia isolate Mad1 (GenBank Accession J02227) cloned into the pBR322 plasmid at the *EcoRI* restriction site (resulting construct: pM1TC) was a generous gift from Walter Atwood (Brown University). JCV stocks were produced by transfection and infection of cells similar to what has been previously described (Hara et al., 1998). Briefly, the viral genome was first extracted from the plasmid backbone by digestion with *EcoRI* (NEB) and re-circularized using T4 DNA ligase.. The resulting viral genomes were then purified using the QIAquick PCR Purification Kit (Qiagen) to prepare for transfection into cells. COS-7 cells, a cell line supportive of JCV replication (Hara et al., 1998), were seeded 1×10^6^ cells per T75 flask and transfected with 2-4 µg of viral genomes using Lipofectamine-2000 (Invitrogen) and Opti-MEM (Gibco) according to manufacturer’s protocol. Transfected cells were incubated at 37°C with 5% CO_2_ for 4 hours, then transfection medium was replaced with infection medium (DMEM supplemented with 2% FBS and 1X Pen/Strep). Cells were cultured at 37°C with 5% CO_2_ for 6-10 weeks, until cytopathic effects (CPE) became evident. During this time, 2-3 mL fresh infection media was added every 3-4 days and cells were split by 1:2 to 1:3 dilution factors once a week to maintain cell health and prevent overcrowding. Upon observation of significant CPE, cells were collected by scraping, combined with culture media, subjected to three freeze-thaw cycles to release intracellular virus. These resulting viral stocks were titrated and stored at −80°C.

### Virus purification

Purified BKV was prepared as previously described (Jiang et al., 2009). Briefly, crude lysate containing high-titer BKV was cleared by centrifugation at 3200 rpm for 30 minutes at 4°C, and supernatant (S1) was separated from the resulting pellet. The pellet (P1) was resuspended in buffer A (10 mM HEPES, pH 7.9, 1 mM CaCl_2_, 1 mM MgCl_2_, 5 mM KCl). The resuspended pellet pH was lowered to 6.0 with 0.5 M HEPES (pH 5.4), and incubated with neuraminidase (1U/mL; Sigma) for 1 hour at 37°C. Pellet buffer pH was then raised pH 7.4 with 0.5 M HEPES (pH 8), and cleared by centrifugation at 16,000 x g for 5 minutes at 4°C. The resulting supernatant (S2) was pooled with the initial (S1), and the pellet (P2) was resuspended in buffer A containing 0.1% deoxycholate (Sigma), incubated for 15 minutes at room temperature, cleared by centrifugation at 16,000xg for 5 minutes at 4°C, and the resulting supernatant (S3) was pooled with the other supernatant fractions. Pooled supernatants were placed over a 4 mL 20% (w/v) sucrose solution and centrifuged at 83,000 x g for 2 hours at 4°C in a SW32Ti rotor (Beckman). The resulting pellet was resupended in 2 mL buffer A, and placed over a CsCl gradient from 1.2-1.4g/cm^3^ in buffer A generated using a J17 gradient former (Jule, Inc.), and centrifuged at 35,000 rpm for 16 hours at 4°C in an SW41 rotor (Beckman). The BKV band formed in the gradient was collected using an 18-guage needle, and dialyzed in a Slide-A-Lyzer Dialysis Cassette, 10K MWCO (ThermoFisher Scientific) over 2 days in 2L buffer A at 4°C, with buffer exchanged once during dialysis. BKV was then aliquoted and stored at −80°C.

### Antibodies and reagents

The following primary antibodies were used in this study: monoclonal mouse anti-SV40 T-antigen (PAb416, EMD Millipore;) at 1:200 for immunofluorescent staining (IF), monoclonal mouse anti-BKV VP1 antibody (in-house generated) at 1:500 for IF, polyclonal rabbit anti-SV40 VP1 (Abcam) at 1:500 for IF and1:1000 for immunoblotting (IB), polyclonal rabbit anti-SV40 VP2/3 (Abcam) at 1:1000 for IF and 1:1000 for IB, polyclonal rabbit anti-BiP (Abcam) at 1:750 for IF and 1:1000 for IB, and monoclonal mouse anti-HSP90 (Abcam) at 1:1000 for IB. The following secondary antibodies were used in this study: in IF applications, goat anti-mouse IgG conjugated to either Alexa Fluor 488, 594, or 647 (ThermoFisher Scientific), goat anti-rabbit IgG conjugated to either Alexa Fluor 488 or 594 (ThermoFisher Scientific); in IB applications, goat anti-mouse IgG conjugated to IRDye 680RD (Li-COR), goat anti-rabbit IgG conjugated to IRDye 800CW (Li-COR). The human anti-BKV VP1 IgG1 antibody P8D11 was produced by the Novartis Institutes for BioMedical Research Biologics Center. D1_min_, D1_min_ W293A, D1_min_ L297A, D1_min_ Y302A, and biotin-peptide probe for the biochemical assay were synthesized and HPLC-purified by the Tufts University Core Facility with purity ≥ 90%. TAT-tagged D1_min_ peptides were synthesized by CPC Scientific. All other peptides were synthesized by the Sigma Chemical Company. Purity was determined by LCMS to be 35-74%.

### Immunofluorescent staining

For T-antigen staining, cells were fixed with 4% paraformaldehyde (w/v) in PBS for 15 minutes, then incubated with primary antibody in 0.2% gelatin, 0.1% Triton X-100 in PBS for 1 hour, followed by incubated with secondary antibody at 1:3000 and 4′,6-diamidino-2-phenylindole (DAPI, Calbiochem) contrast stain at 1.67 μg/mL in 0.2% gelatin in PBS for 1 hour. For VP1 co-localization and cell-binding assays, cells were fixed with 4% paraformaldehyde (w/v) in PBS for 15 minutes then permeabilized with 0.1% Triton X-100 in PBS for 10 minutes. Cells were then blocked with 2% goat serum (Invitrogen) for 30 minutes, then incubated with primary antibodies for 1 hour, and secondary antibodies for 1 hour, followed by a 10 minute incubation with DAPI contrast stain at 1.67 μg/mL (Calbiochem). For VP2/3 staining, cells were fixed in 100% methanol for 15 minutes at −20°C then blocked in 3% nonfat milk (Bio-Rad), 0.1% Tween-20 (Bio-Rad) in PBS for 30 minutes. Cells were then incubated with anti-VP1 and anti-VP2/3 antibodies for 1 hour, and secondary antibodies for 1 hour, followed by a 10 minute incubation with DAPI contrast stain at 1.67 μg/mL.

### Infections

Viral titers were measured by fluorescent focus assay, as previously described (Jiang et al., 2009). For 96-well plate format assays, RPTE cells were seeded 12,000 per well. For ER-to-cytosol retrotranslocation assays, RPTE cells were seeded in 6-well plates at 380,000 cells per well. For non-synchronized infections of RPTE cells, virus was diluted in RenaLife medium and added to cells followed by incubation at 37°C for the desired time. For synchronized infections, cells were pre-chilled to 4°C for 15 minutes. Purified virus was diluted into cold RenaLife medium and incubated with cells for 1 hour at 4°C. Cells were rinsed once with cold RenaLife medium, followed by addition of warm medium and incubation at 37°C for the desired time. For COS-7 cell assays, cells were seeded 5,000 per well in a 96-well plate format. COS-7 cells were infected using a synchronized infection protocol as described above, with the following modifications: JCV or BKV were diluted into low-serum medium (DMEM supplemented with 2% FBS), and cells were rinsed with cold low-serum medium and cultured in low-serum medium at 37°C for the desired time.

Spreading infection assays were performed as follows. Re-circularized BKV genomes were prepared as described. RPTE cells were reverse-transfected with 100 ng viral genome DNA using Lipofectamine 3000 (Invitrogen) at a 1.5:1 ratio of L3000 to DNA, and Opti-MEM (Gibco) in a 96 well-plate format. Medium was exchanged the following day, and plates were incubated at 37°C for the desired time.

### Preparation of BKV mutants

Mutant BKV genomes were prepared using PCR site-directed mutagenesis using the primers listed below.

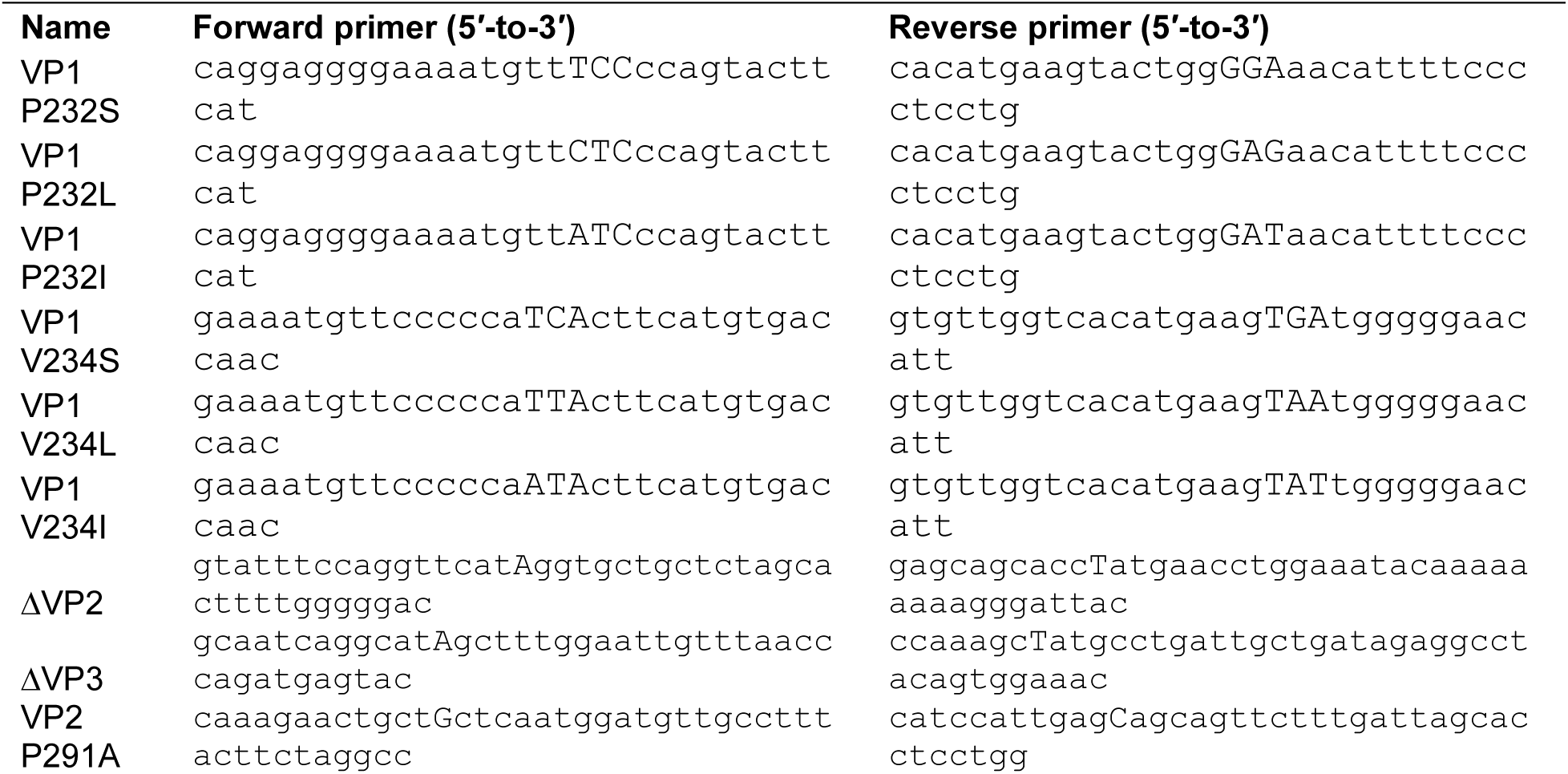
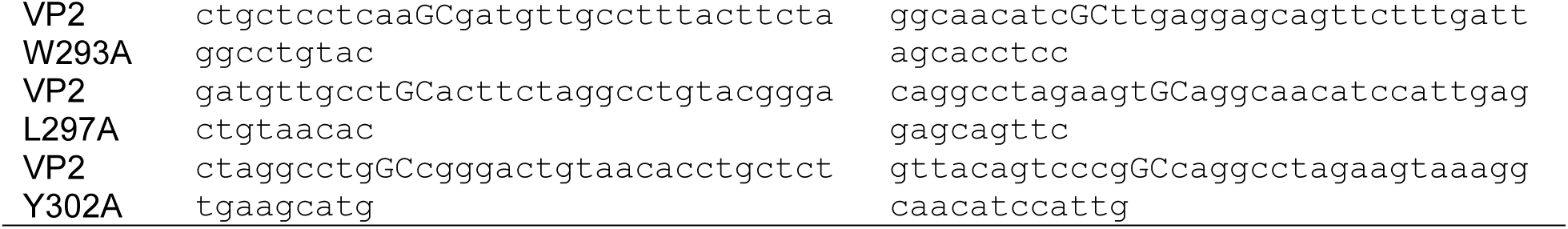

After PCR, reactions were treated with *DpnI* (NEB) to remove template DNA, and PCR products were used to transform XL10-Gold (VP1 mutants, Agilent) or 10-beta cells (VP2 mutants, NEB). Resultant colonies were sequenced and analyzed for desired mutation, and viral genomes were prepared as described. Point mutations lead to amino acid substitutions in VP2 and VP3. Deletion mutants were obtained by point mutation of the start codon. While ΔVP2 did not affect VP3 sequence, ΔVP3 resulted in a M120I substitution in VP2. BKV ΔVP2ΔVP3 genome was generated by successive rounds of site-directed mutagenesis with ΔVP2 and ΔVP3 primer sets.

### Inhibitor treatments

Dose-response curves were determined using 3-fold, 10-point titrations of inhibitor. For peptide EC_50_ determination, cells were treated with inhibitors for two hours prior to infection. For synchronized infection treatments, inhibitors were added immediately following synchronization; EC_50_ concentration for P8D11 was determined using the synchronized treatment protocol. CC_50_ values were determined using a CellTiter-Glo luminescent cell viability assay (Promega) after two days of treatment, with luminescence detected on a PHERAstar FS (BMG Labtech).

### Cell-binding assay

Purified infectious BKV virions were incubated with titrated concentrations of D1_min_ peptide or P8D11 antibody for 1 hour at 4°C in RenaLife medium. RPTE cells seeded in 96-well plate format were cooled for 15 minutes at 4°C, and medium containing virus and inhibitor mix was added to cells and incubated for 1 hour at 4°C. Cells were rinsed with cold RenaLife medium, and proceeded to fixation and staining as described.

### Time of addition assay

RPTE cells seeded in 96-well plate format were subjected to a synchronized infection at low MOI (MOI = 0.3). Timepoint 0 h.p.i. samples were treated with inhibitor compounds immediately after synchronized infection, others added at the timepoint indicated. Inhibitors were used at the following concentrations: D1_min_, D1_min_-TAT, TAT-D1_min_, 0.8 μM; P8D11, 0.014 μM (2 μg/mL). Plates were incubated at 37°C and fixed at 48 h.p.i.

### Fractionation assay

RPTE cells were cultured in 2 wells in 6-well format per treatment. Cells were subjected to synchronized infection at high MOI (MOI = 10), and treated with 5 μM peptide immediately after synchronized infection. Cells were harvested 24 h.p.i. with 0.05% trypsin, 0.02% EDTA (Lifeline Cell Technology) for 2 minutes until cells were detached. Trypsin was inhibited with an equal volume of Trypsin Neutralizing Solution (Lifeline Cell Technology), and wells rinsed with phosphate-buffered saline (PBS). Cells were pelleted at 90 x g for 5 minutes at 4°C, and rinsed with 1 mL cold PBS buffer. Cell pellets were then lysed in 50 μl HNF buffer (150 mM HEPES pH 7.2, 50 mM NaCl, 2 mM CaCl_2_) containing 0.025% digitonin (Thermo Lifesciences) and 1X cOmplete^TM^, Mini Protease Inhibitor Cocktail (Roche) for 10 minutes on ice. Lysates were then clarified with a 21,100 x g centrifugation at 4°C for 10 minutes. The supernatant (cytosolic fraction) was removed and the pellet was rinsed with 1 mL HNF buffer and transferred to a fresh tube, and pelleted again at 21,100 x g centrifugation at 4°C for 10 minutes. Pellets were then resuspended in sample buffer directly in sample buffer. Samples were boiled at 95°C for 10 minutes then stored at −20°C until subjected to SDS-PAGE and immunoblotting.

### Immunoblotting

Samples were prepared in sample buffer (NuPAGE LDS Sample Buffer (Invitrogen), 100 mM dithiothreitol (DTT)), boiled for 5 minutes at 95°C, and subjected to SDS-PAGE using 4-12% Bis-Tris Bolt gels (Invitrogen) in MOPS running buffer (Invitrogen). Proteins were transferred to Immobilon-FL PVDF membrane (Millipore) in transfer buffer containing 1X NuPAGE transfer buffer (Invitrogen), 20% methanol, and 0.05% SDS for 70 minutes. Membranes were blocked using Odyssey blocking buffer (TBS) (Li-COR) for 30 minutes at room temperature, and incubated with primary antibody diluted in Odyssey blocking buffer containing 0.05% Tween 20 overnight at 4°C. Membranes were rinsed three times with Tris-buffered saline (TBS) containing 50 mM Tris-HCl, pH 7.4, 150 mM NaCl, and 0.1% Tween 20 (TBS-T), and incubated with secondary antibodies for 1 hour at room temperature. Membranes were again rinsed three times with TBS-T and once with TBS containing no detergent, and membranes were immediately scanned using an Odyssey Infrared Imager (Li-COR).

### Imaging and image segmentation

Images were acquired on an ImageXpress Micro XLS system (Molecular Devices) with a 10x objective for assays determining percent infected cells (Nikon CFI Plan Fluor, MRH00101), or a 60x objective for all other assays (Nikon 60X Plan Apo λ, MRD00605). Images were processed using CellProfiler version 2.1.2 (Kamentsky et al., 2011).

### Data analysis

EC_50_ values were calculated using XLFit v5.5.0.5 (IDBS). Quantification and processing of data generated by CellProfiler was performed using R v3.5.1 (R Core Team, 2018). Microscopy data was quantified per field of view, averaged per well, and data displayed as the mean value ± SD across replicate wells. Immunoblot images were processed using Fiji built on ImageJ v1.52b (Schindelin et al., 2012).

### VP1 plasmid construction

Synthetic DNA, codon optimized for Sf9 cell expression, encoding full-length BKV serotype 1 VP1 and JCV VP1 were generated for expression of VP1 proteins. For VLP production, DNA fragments encoding full length VP1 were inserted into the pFastBac1 plasmid and baculovirus was generated following the Bac-to-Bac method (Invitrogen). For VP1 pentamer production, DNA fragments encoding either BKV VP1 residues (2-362), JCV VP1 residues (2-354), or BKV VP1 residues (30-297) were inserted into a gateway adapted pGEX plasmid for expression in E. coli with an N-terminal GST-6xHis-Tev tag. The mutations P232S, V234S, and T224A/T243A were introduced into the BKV VP1_30-297_ plasmid by QuikChange site-directed mutagenesis.

### VP1 pentamer production

BL21 Star (DE3) E. coli cells were transformed with expression plasmids and plated on LB agar plates supplemented with 100 µg/ml carbenicillin. Cell were grown in Terrific broth (supplemented with 15 mM sodium phosphate pH 7.0, 2 mM MgCl_2_, 100 µg/ml carbenicillin) with shaking at 37°C until the OD600 reached 0.7, the temperature was then lowered to 18°C, and Isopropyl β-D-1 thiogalactopyranoside (IPTG) added to 0.5 mM. After 16 hours, harvested cells by centrifugation and stored at −80°C. Cells were re-suspended in chilled lysis buffer (25 mM Tris-HCl pH 8.0, 200 mM NaCl, 5% glycerol, 1 mM Tris(2-carboxyethyl) phosphine (TCEP), 15 mM Imidazole, 1X Roche complete EDTA-free protease inhibitor cocktail, 1X Pierce universal nuclease) at a ratio of 5 mL buffer per gram of cell paste. Cells were lysed by passing through an M-110P microfluidizer at 17,500 PSI, on ice. The lysate was centrifuged at 26,800 x g for 45 minutes at 4°C. Equilibrated 10 mL Nickel sepharose Fast Flow (GE) column in lysis buffer and loaded clarified lysate. Washed resin with 3 column volumes (CV) of lysis buffer. Washed resin with 5 CV of wash buffer (25 mM Tris-HCl pH 8.0, 200 mM NaCl, 1 mM TCEP, 40 mM Imidazole, 5% glycerol) and VP1 was eluted with 2 CV elution buffer (25 mM Tris-HCl pH 8.0, 200 mM NaCl, 1 mM TCEP, 250 mM Imidazole, 5% glycerol). The N-terminal tag was removed by cleavage with Tev protease and pentamers were loaded onto a Superdex 200 column equilibrated in SEC buffer (25 mM Tris-HCl pH 8.0, 100 mM NaCl, 1 mM TCEP, 5% glycerol). Pooled peak fractions and concentrated using a 50,000 MWCO Amicon concentrator.

### Labeled VP1 pentamer production

BL21 Star (DE3) E.coli cells transformed with VP1 expression plasmid were grown in M9 minimal media (6 g/L Na_2_HPO_4_, 3 g/L KH_2_PO_4_, 0.5 g/L NaCl, 2 mM MgSO_4_, 0.1X Vitamins (Sigma R7256), 1 g/L ^15^NH_4_Cl, 3 g/L glucose, 100 µg/mL carbenicillin, 0.1X trace metals) made up in D_2_O. Cells were incubated at 37°C with shaking at 250 RPM until the OD600 reached ∼0.7. To each liter of culture added 70 mg of 2-Ketobutyric acid-4-^13^C,3,3-d2 sodium salt hydrate, 120 mg of 2-Keto-3-methyl-^13^C-butyric-4-^13^C, 3-d acid sodium salt, 100 mg deuterated glycine, and 100 mg L-threonine (4-^13^C;2,3-D2). After 1 hour, the temperature was lowered to 24°C and IPTG was added to 250 µM. After 16 hours, cells were harvested by centrifugation and stored at −80°C. VP1 pentamers were purified as described above with the exception of the SEC buffer being 50 mM Na-Phosphate pH 7.0, 100 mM NaCl.

### VLP production

Recombinant baculovirus encoding untagged full length BKV VP1 or JCV VP1 were used to infect Sf9 insect cells in suspension at 1.5 million cells/mL, the cells were incubated at 27°C with shaking at 120 RPM for 72 hours then harvested by centrifugation and stored at −80°C. Cells were re-suspended in lysis buffer (20 mM Tris-HCl pH 7.5, 1 M NaCl, 1X Roche EDTA free protease inhibitor cocktail) at a ratio of 10 mL lysis buffer per gram of cell pellet and lysed by sonication on ice then the lysate was centrifuged at 16,000 x g for 20 minutes at 4°C. The supernatant was layered onto 3ml of 40% glucose made up in 1X PBS and centrifuged at 116,000 x g for 2.5 hours at 4°C. Dissolved pellet in IEX buffer A (25 mM Tris-HCl pH 8.0, 25 mM NaCl) and loaded onto a 10 mL Sepharose Q-HP (GE) column equilibrated in IEX buffer A. Washed column with 3CV IEX buffer A and eluted with a linear NaCl gradient from 25 mM to 700 mM NaCl across 25 CV. Pooled peak fractions and loaded onto 10 mL Capto Core 700 (GE) resin equilibrated in SEC buffer (25 mM Tris-HCl pH 8.0, 100 mM NaCl) collecting the flow-through fraction. Loaded onto a Sephacryl S500 26/60 column (GE) and collected peak fractions, concentrated with a 100,000 MWCO Amicon concentrator.

### Protein reagents

The following VP1 sequence was used for the biochemical AlphaScreen assays:

> BKV serotype 1 VP1 (30-297)

GKGGVEVLEVKTGVDAITEVECFLNPEMGDPDENLRGFSLKLSAENDFSSDSPERKMLPCYSTARIPLPNLNEDLTCGNLLMWEAVTVQTEVIGITSMLNLHAGSQKVHEHGGGKPIQGSNFHFFAVGGDPLEMQGVLMNYRTKYPEGTITPKNPTAQSQVMNTDHKAYLDKNNAYPVECWIPDPSRNENTRYFGTFTGGENVPPVLHVTNTATTVLLDEQGVGPLCKADSLYVSAADICGLFTNSSGTQQWRGLARYFKIRLRKRSVK

The following VP1 sequence was used for SPR assays:

>BKV serotype 1 VP1 (2-362)

GAPTKRKGECPGAAPKKPKEPVQVPKLLIKGGVEVLEVKTGVDAITEVECFLNPEMGDPDENLRGFSLKLSAENDFSSDSPERKMLPCYSTARIPLPNLNEDLTCGNLLMWEAVTVQTEVIGITSMLNLHAGSQKVHEHGGGKPIQGSNFHFFAVGGDPLEMQGVLMNYRTKYPEGTITPKNPTAQSQVMNTDHKAYLDKNNAYPVECWIPDPSRNENTRYFGTFTGGENVPPVLHVTNTATTVLLDEQGVGPLCKADSLYVSAADICGLFTNSSGTQQWRGLARYFKIRLRKRSVKNPYPISFLLSDLINRRTQRVDGQPMYGMESQVEEVRVFDGTERLPGDPDMIRYIDKQGQLQTKML

### Chemical attachment of biotin to VP1 proteins

For SPR analysis, biotin was covalently attached to VP1(2-362) with the sulfo-N-hydroxysuccinimide (NHS) ester of a biotin derivative (ThermoFisher Scientific # 21338) as follows: To a 1500 μl of a solution of BKV VP1 (2-362) protein at 17 μM in PBS buffer containing 1 mM TCEP was added 8 μl of a 1 mg/mL (1.5 mM) solution of sulfo-NHS-LC-LC-biotin in water. The solution was mixed with a vortex mixer briefly (1 second), and incubated at room temperature for one hour. The solution was transferred to a ThermoFisher Slide-A-Lyzer dialysis cassette (3.5 kDa molecular weight cut-off, 3 mL) and dialyzed extensively against 3 times 2 liters of PBS buffer containing 1 mM TCEP at 4°C for 18 hours.

### Analysis of peptide:VP1 interactions by surface plasmon resonance

SPR analysis for the determination of the dissociation constant *K*_D_ was performed with a Biacore T200 instrument with PBS buffer containing 1 mM TCEP, 0.05% Tween-20 (or P20, GE Healthcare) and 1 mM ethylenediaminetetraacetic acid (EDTA) at 20°C. The flow rate was 60 μl per minute. VP1_2-362_ protein covalently modified with biotin was loaded onto a streptavidin-coated Biacore biosensor (GE Healthcare Series S Sensor Chip SA, catalog BR-1005-31) that had been pre-treated with 50 mM sodium hydroxide containing 1 M sodium chloride. The protein loading response was 6000-8000 resonance units. Peptides were analyzed using the single cycle kinetics method according to instrument control software instructions. Data were analyzed using Biacore Evaluation Software to generate affinity constants (*K*_D_).

### AlphaScreen competitive binding assay

The assay was run in a Tris buffer at pH 7.5 containing 100 mM NaCl, 0.01% Tween 20, 1 mM EDTA and 0.01% bovine serum albumin. The D1_22_ biotin-peptide probe with the sequence [H]-APGGANQRTAPQWMLPLLLGLYG-GGGK(Biotin)-[OH] was incubated with BKV VP1_30-297_ for 90 minutes before addition of an anti-BKV VP1 antibody (in-house generated) along with AlphaScreen streptavidin donor and protein A acceptor beads (PerkinElmer). Samples were incubated overnight before reading on a PerkinElmer Envision. Untagged peptides were assessed in a competition mode where they were serially diluted in assay buffer and added to the VP1 along with the biotin-peptide probe. Peptide IC_50_ values (n=3) were calculated in Microsoft Excel using XLfit.

### Co-crystallization of BKV VP1 pentamer with D1_min_

For co-crystallization, 13-mer D1_min_ peptide was added to the protein to a final concentration to ratio of 5:1 D1_min_ to pentamer. The resulting mixture was incubated on ice for 1 hour and was concentrated to 15 mg/mL protein overall. Prior to crystallization, the mixture was passed through a 0.2-micron filter. The protein-peptide complex was crystallized using the hanging drop vapor diffusion method. 2.0 µL of protein solution was mixed with 2.0 µL of well solution, which consisted of 20% PEG-3350, 5% ethylene glycol, 0.1 M Tris buffer pH 8.5, 10 mM TCEP. The resulting drop was suspended over a reservoir of 0.3 mL well solution. The crystals grew at 18°C for approximately 12-24 hours. Crystals were washed briefly in a cryoprotectant consisting of 80% well and 20% ethylene glycol (v/v) and then flash-frozen in liquid nitrogen prior to data collection.

### Structure solution and refinement of BKV VP1 pentamer:D1_min_ complex

The X-ray diffraction data was collected at a wavelength of 1.54187Å and a temperature of 100°K on a Rigaku FRE+ anode utilizing a Decris 300K Pilatus detector. Data integration and scaling were performed by using the autoPROC implementation of XDS and AIMLESS (Vonrhein et al., 2011). The structure of the complex was solved via Molecular Replacement using the CCP4i Suite implementation of PHASER (McCoy et al., 2007; Winn et al., 2011). The structure was built and refined via alternating rounds of real-space rebuilding in Coot and refined using autoBuster (Global Phasing) until convergence was reached. Data reduction and structure refinement statistics are presented in **Supplemental Table S3**. After attempts to refine the D1_min_ model to convergence, with suitable φ/ψ angles as defined by the Ramachandran plot and suitable rotamers, it became clear that a single binding model could not account for the electron density seen in 2Fo-Fc maps. A model that rationalized the electron density was achieved by fitting the peptide with an occupancy of 0.8. Subsequent reciprocal space and real-space refinement of this model led to a suitable fit of the peptide into the observed density and reduced, but still unaccounted for, difference density for the remaining symmetry-related binding modes, which are not fit. As such, the co-structure is presented as a model constructed using the observed density, or a “structurally-guided model.” Alignment of structurally-guided model with apo BKV pentamer X-ray structure (PDB: 4MJ1; Neu et al., 2013) was performed using the align tool in PyMOLv2.2.3 (Schrödinger, LLC) with a 10Å cutoff for outliers. Post-alignment, RMSD values were calculated using the tool rms_cur without refitting the alignment.

### NMR spectroscopy

All NMR experiments were performed on a Bruker Avance III 600 MHz spectrometer equipped with a 5 mm z-gradient QCI-F cryo probe. The temperature in all experiments was 32°C (305K). The NMR samples were prepared in 160 μL PBS buffer at pH 7.5, containing 2 mM deuterated DTT, 10% (v/v) D_2_O and 11.1 μM 4,4-dimethyl-4-silapentane-1-sulfonic acid (DSS, internal standard). The final protein concentration of truncated (aa 30-297), ^2^H,^12^C,^15^N and ^1^H,^13^C-methyl-ILVT labeled BKV VP1 was 125 μM (monomer concentration) in all experiments. The 13-mer D1_min_ (Ac-APQWMLPLLLGLY-NH_2_) and alanine substitution peptide D1_min_ W293A (Ac-APQAMLPLLLGLY-NH_2_) were dissolved in d_6_-DMSO and added to the protein at various concentrations (6.25 μM – 200 μM).

1D-^1^H NMR experiments were acquired with 64 scans, excitation sculpting water suppression and a relaxation delay of 2 seconds. 2D ^1^H,^13^C-HMQC SOFAST (Schanda et al., 2005) spectra were recorded using 50% non-uniform sampling, 1024 and 256 points in the direct and indirect dimensions, respectively, 192 scans and a recycling delay of 200 milliseconds. Spectra were processed and analyzed using TOPSPIN version 3.5. Methyl peak assignments were obtained by means of twelve amino acid point mutations (L68A, L254M, V136I, V231I, V234I, T46S, T118S, T224S, T238S, T240S, T277S, I45V; see **Supplement Figure S3B** for an example of this method) and by using a methyl walk approach based on ^13^C-resolved 4D-HMQC-NOESY-HMCQ experiments (Proudfoot et al., 2016).

## ACKNOWLEDGEMENTS

We thank Weidong Zhong, Kelly Wong, Dirksen Bussiere, and Don Ganem for their leadership roles, Catherine Jones for important intellectual contributions in project team management, Atul Sathe, Lihong Zhao, and Sue Ma for their guidance, advice, and assistance with polyomavirus virology, and members of the local postdoctoral community for their support. All authors were funded by Novartis Institutes for Biomedical Research. J.R.K is a postdoctoral fellow at Novartis Institutes for Biomedical Research. S.F., J.S., A.F., A.O.F., M.K., P.K., E.O., C.C., J.R.A, and C.A.W. are or were employees of Novartis Institutes for Biomedical Research at the time of these studies.

## AUTHOR CONTRIBUTIONS

J.R.K, J.R.A., and C.A.W. wrote the manuscript. J.R.K performed peptide antiviral assays, peptide cytotoxicity assays, *in vitro* peptide affinity purifications, fractionation assays, VP2/3 exposure studies, and associated data analysis. S.F. and C.A.W. performed biophysical and biochemical assays. J.S. performed purification of protein reagents and VLPs. M.K., D.B., E.O., and C.C. performed XRC studies. A.F. and A.O.F. performed NMR experiments. J.R.K. and J.R.A. performed time of addition study. J.R.K. and P.K. performed mutant virus spread assays.

## SUPPLEMENTAL FIGURES

**Supplemental Figure S1**

**A.** Multiple sequence alignment of VP1 protein from BKV (P03088), JCV (P03089), SV40 (P03087), murine polyomavirus (MPy; P03090). VP1 pore residues that are proximal to D1_min_ peptide binding region are highlighted. Dots indicate conserved sequence, relative to BKV. **B.** Multiple sequence alignment of VP2 proteins (BKV: P03094, JCV: P03095, SV40: P03093, MPy: P03096). Sequence from which D1_min_ is derived is highlighted. Alignment was performed using Clustal Omega (Madeira et al., 2019).

**Supplemental Figure S2**

**A.** ^1^H,^13^C-HMQC spectra showing peptide (100 µM) induced perturbations of tr-VP1 (125 µM protein + 25 µM peptide; black) ILVT methyl signals. Left: the wild-type peptide (magenta) induces secondary CSPs (e.g. I45, see inset in upper corner), indicating micromolar binding to a second site. The affected residues are clustered in the lower pore. Right: in contrast to the wild-type D1_min_, the W293A alanine-substituted peptide (red) hardly induces secondary chemical shifts. This observation suggests that secondary binding is very weak (millimolar *K*_D_). **B.** ^1^H,^13^C-HMQC spectra showing an inset of tr-VP1-I45V (100 µM; red) and wild-type tr-VP1 (100 µM; black) Ile methyl signals. The isoleucine signal at 0.84 ppm (^1^H) / 12.80 ppm (^13^C) disappears when I45 gets mutated, making it possible to assign the residue. **C.** ^1^H,^13^C-HMQC spectra showing peptide (200 µM) induced perturbations of tr-VP1 (125 µM protein + 25 µM peptide; black) ILVT methyl signals. Left: most of the VP1 peaks affected by the second site binding of the wild-type peptide (magenta) show severe line broadening at 200 µM ligand concentration (see inset in upper corner). This effect may be caused by specific or unspecific binding of multiple copies of the peptide to the lower pore. Right: the W293A alanine-substituted peptide (red) does not induce super-stoichiometric line broadening as its affinity for the secondary binding site is very low. **D-E.** Highlighting primary binding site (green) and secondary binding site (yellow) VP1 residues that show peptide induced CSPs (PDB: 4MJ1; Neu et al., 2013).

**Supplemental Figure S3**

**A.** Overlay of X-ray crystal structures from wild-type (magenta) or P232S (cyan) VP1 pentamer.

**B.** Overlay of X-ray crystal structures from wild-type (magenta) or V234S (bronze) VP1 pentamer.

**C.** Cutaway view of structurally guided model of D1_min_ peptide bound to BKV VP1 showing 3 VP1 molecules within the pentamer with D1_min_ peptide bound. 2Fo-Fc electron density map, contoured at 1σ, is shown in blue. Experimental electron density is observed to occupy the upper region of the pore and has a distinct helical appearance. **D.** Alignment of structurally-guided model of ligand-bound (peptide not displayed) BKV pentamer (blue) to apo BKV pentamer (orange) (PDB: 4MJ1; Neu et al., 2013) (RMSD: 0.85Å), with magnified view of pore residues T224, T226, V231, P232, and V234 (RMSD: 0.42Å, using previous whole-pentamer alignment).

**Supplemental Figure S4**

**A.** Coomassie-stained SDS-PAGE of streptavidin-based affinity purification of biotinylated D1_min_ peptides, assaying for co-affinity purification of VP1 protein in Figure 4. Sequences for peptides used in this assay can be found in **Supplemental Table S2**. VLP: virus-like particle.

**Supplemental Figure S5**

**A.** Immunoblot for VP1 protein expressed from transient transfection of constructs used in spreading infection assay for **Figure 4E**. **B.** Immunoblot for VP2/3 protein expressed from transient transfection of constructs used in spreading infection assay for **Figure 4F**.

## SUPPLEMENTAL TABLES

**Supplemental Table S1.**
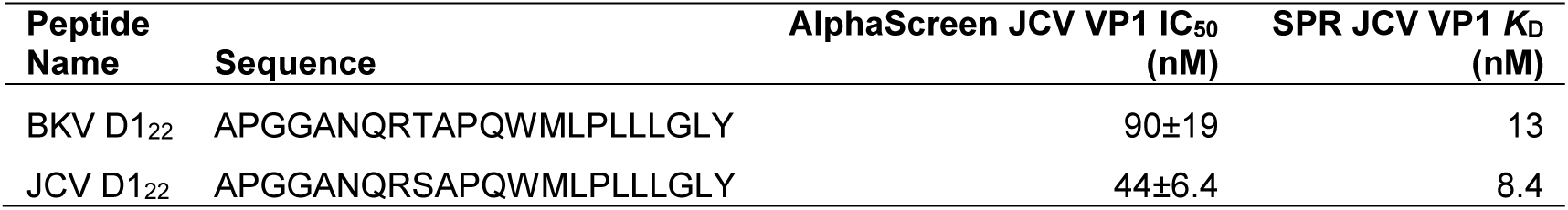
JCV VP1 binding data

**Supplemental Table S2.**
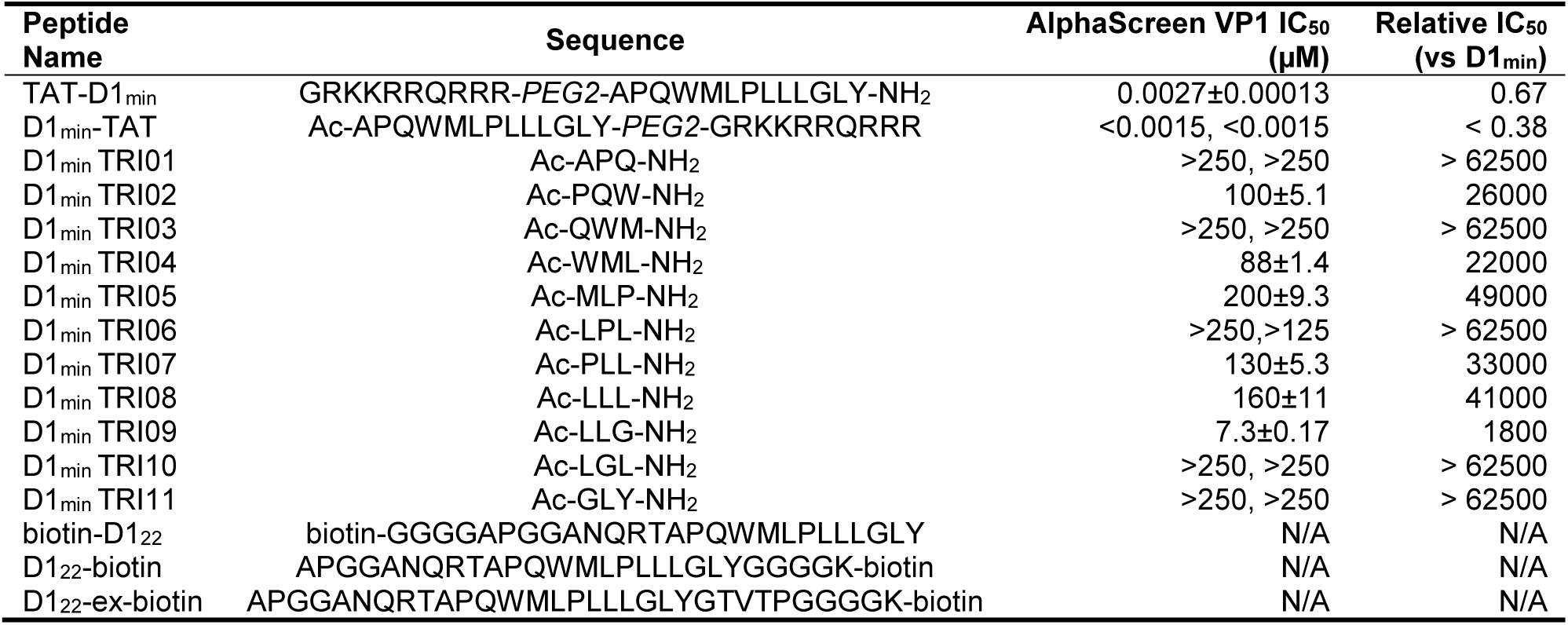
Additional peptide information

**Supplemental Table S3.** Data collection and refinement statistics (molecular replacement)

| BKV VP1 pentamer-D1<sub>min</sub> |  |
| --- | --- |
| **Data collection** |  |
| Space group | P 21 21 21 |
| Cell dimensions |  |
| *a*, *b*, *c* (Å) | 89.47, 124.27, 127.83 |
| α, β, γ (°) | 90.00, 90.00, 90.00 |
| Resolution (Å) | 89.10 – 2.36 |
| *R*<sub>sym</sub> or *R*<sub>merge</sub> | 0.09(0.705) |
| *CC*<sub>1/2</sub> | 0.972(0.30) |
| *I* / *σI* | 14.9(4.5) |
| Completeness (%) | 99.9 (99.7) |
| Redundancy | 22.0(14.7) |
| **Refinement** |  |
| Resolution (Å) | 44.70-2.36 |
| No. reflections | 16198 |
| *R*<sub>work</sub> / *R*<sub>free</sub> | 0.199/0.257 |
| No. atoms |  |
| Protein | 3106 |
| Ligand/ion | 2 |
| Water | 195 |
| *B*-factors |  |
| Protein | 58.63 |
| Ligand/ion | 34.10 |
| Water | 50.32 |
| R.m.s. deviations |  |
| Bond lengths (Å) | 0.010 |
| Bond angles (°) | 1.13 |

**Supplemental Table S4.** Control inhibitor half-maximal effective concentrations (EC_50_) and half-maximal cytotoxic concentrations (CC_50_).

| Inhibitor | Type | EC<sub>50</sub> (nM) | CC<sub>50</sub> (μM) |
| --- | --- | --- | --- |
| TAT-D1<sub>min</sub> | peptide | 100 | 9.6 |
| D1<sub>min</sub>-TAT | peptide | 79 | 7.5 |
| P8D11 | monoclonal antibody | 0.35<br>(50 ng/mL) | Not observed |

