## Supplemental Figure S1 for "A VP2/3-derived peptide exhibits potent antiviral activity against BK and JC polyomaviruses by targeting a novel VP1 binding site"

A

```
BKV     1 MAPTKRKG--ECPGAAPKKPKEPVQVPKLLIKGGVEVLEVKTGVDAITEVECFLNPEMGDPDENL--------RGFS  67
JCV     1 ........--.--------R.D.........R.............S........T........H.--------....  59
SV40    1 ........--S..................V....I...G......SF.........Q..N...HQ--------K.L.  67
MPy     1 ...KRKS.VSK.ETKCT.ACPR.AP.........M...DLV..P.SV..I.A....R..Q.PTPESLTEGGQYY.W.  77

BKV    68 LKLSAENDFSSDSPERKMLPCYSTARIPLPNLNEDLTCGNLLMWEAVTVQTEVIGITSMLNLHAGSQ---KVHEHGG 141
JCV    60 KSI.ISDT.E....N.D......V................I.......LK.....V..LM.V.SNG.---AT.DN.A 133
SV40   68 KS.A..KQ.TD...DKEQ.....V.......I........I........K.....V.A.....S.T.---.T..N.A 141
MPy    78 RGINLA-TSDTWI.RNNT..TW.M.KSSF.C.......DT.Q.....S.K...V.SG.L.DV.GFNKTHRFSKHK.N 153

BKV   142 GKPIQGSNFHFFAVGGEPLEMQGVLMNYRSKYPDGTI------TPKNPTAQSQVMNTDHKAYLDKNNAYPVECWVPD 212
JCV   134 ...V..TS....S....A..L....F...T.......------F...A.V.......E........K.......... 204
SV40  142 ....................L....A...T...AQ.V------....A.VD..Q.......V...D........... 212
MPy   154 ST.VE..QY.V..G.....DL..LVTDA.T..KEEGVVTIKTI.K.DMVNKD..L.PIS..K...DGM....I.H.. 230

BKV   213 PSRNENARYFGTFTGGENVPPVLHVTNTATTVLLDEQGVGPLCKADSLYVSAADICGLFTNSS-GTQQWRGLARYFK 288
JCV   205 .T....T.....L...........I...........F.......G.N..L..V.V..M...R.-.S......S.... 280
SV40  213 ..K...T.....Y...........I...........................V........T.-.....K..P.... 288
MPy   231 .AK...T....NY.....TTA....QF...L.......N.......GEG..L.CV..M.WRVTRNYVSSLEK.FP.... 307

BKV   289 IRLRKRSVKNPYPISFLLSDLINRRTQRVDGQPMYGMESQVEEVRVFDGTERLPGDPDMIRYIDKQGQLQTKML--  362
JCV   281 VQ....R...........T.......P..........DA........E...E.......M..V..Y........--  354
SV40  289 .T.................................I.S........YED..E............EF..TT.R.Q--  362
MPy   308 .T....W......MAS.I.S.F.NMLPQ.Q....E.ENT.......Y....PV......T..V.RF.KTK.VFPGN  383
```

B

```
BKV     1 MGAALALLGDLVASVSEAAAATGFSVAEIAAGEAAAAIEVQIASLATVEGITSTSEAIAAIGLTPQTYAVIAGA  74
JCV     1 .............T......................T...E........................E.....T..  74
SV40    1 .....T.....I.T...........................L..V....L-............I..A....S..  73
MPy     1 ....TI.V...IEGLA.VSTL..L.AEA.LS...L..LDGE.TA.-.L..VM.SET.L.TM.ISEEV.GFVSTV  73

BKV    75 PGAI---AGFAALIQTVSGISSLAQVGYRFFSDWDHKVSTVGLYQQSGMALELFNPDEYYDILFPGVNTFVNNI 145
JCV    75 ...V---......V...T.G.AI..L.....A...........F..PA...Q....ED..........A..... 145
SV40   74 .A..---......L...T.V.AV.......................P...VD.YR..D.........Q...HSV 144
MPy    74 .VFVSRT..AIW.M...Q.A.TISLGIQ.YLHNEE--.P.----VNRN...IPWRDPALL..Y.....Q.AHAL 141

BKV   146 QYLDPRHWGPSLFATISQALWHVIRDDIPSI---TSQELQRRTERFFRDSLARFLEETTWTIVNAPINF----- 211
JCV   146 H............S.....F.NLV...L.AL---....I....QKL.VE...........A...S.A.L----- 211
SV40  145 ..........T..NA....F.R..QN...RL---.....E...Q.YL..............VI...V.W----- 210
MPy   142 NVV--HD..HG.LHSVGRYV.QMVVQETQHRLEGAVR..TV.QTHT.L.G...L..N.R.VVS...QSAIDAIN 213

BKV   212 ---------YNYIQQYYSDLSPIRPSMVRQVAEREGTRVHFGHTY--SIDDADSIEEVTQRMDLR-NQQSVHSG 273
JCV   212 ---------....SD...R...V.........Q....YIS...S.TQ........Q.....L..K-T-PN.Q.. 274
SV40  211 ---------..SL.D...T......T......N...LQIS.....D-N..E....QQ..E.WEAQSQSPN.Q.. 274
MPy   214 RGASSASSG.SSLSD..RQ.GLNP.QRRALFNRI..SMGNG.P.P--------------------AAHIQDE.. 267

BKV   274 EFIEKTIAPGGANQRTAPQWMLPLLLGLYGTVTPALE--AYEDGPNQKKRRVSRGSSQKAKGTRASAKTTNKRR 345
JCV   275 ....RS.........S.....................--.......K....K--------E.P...S..SY... 338
SV40  275 .....FE......................S..S..K--.......K...KL.......T...S....ARH... 346
MPy   268 .V.KFYQ.QVVSH..VT.D.....I.....DI..TWATVIE....QK....L---------------------- 319

BKV   346 SRSSRS                                                                     351
JCV   339 ......                                                                     344
SV40  347 N.....                                                                     352
MPy       ------
```
