## Supplementary figures and images for "A VP2/3-derived peptide exhibits potent antiviral activity against BK and JC polyomaviruses by targeting a novel VP1 binding site"

### Supplemental Figure S2

#### Supplemental Figure S2

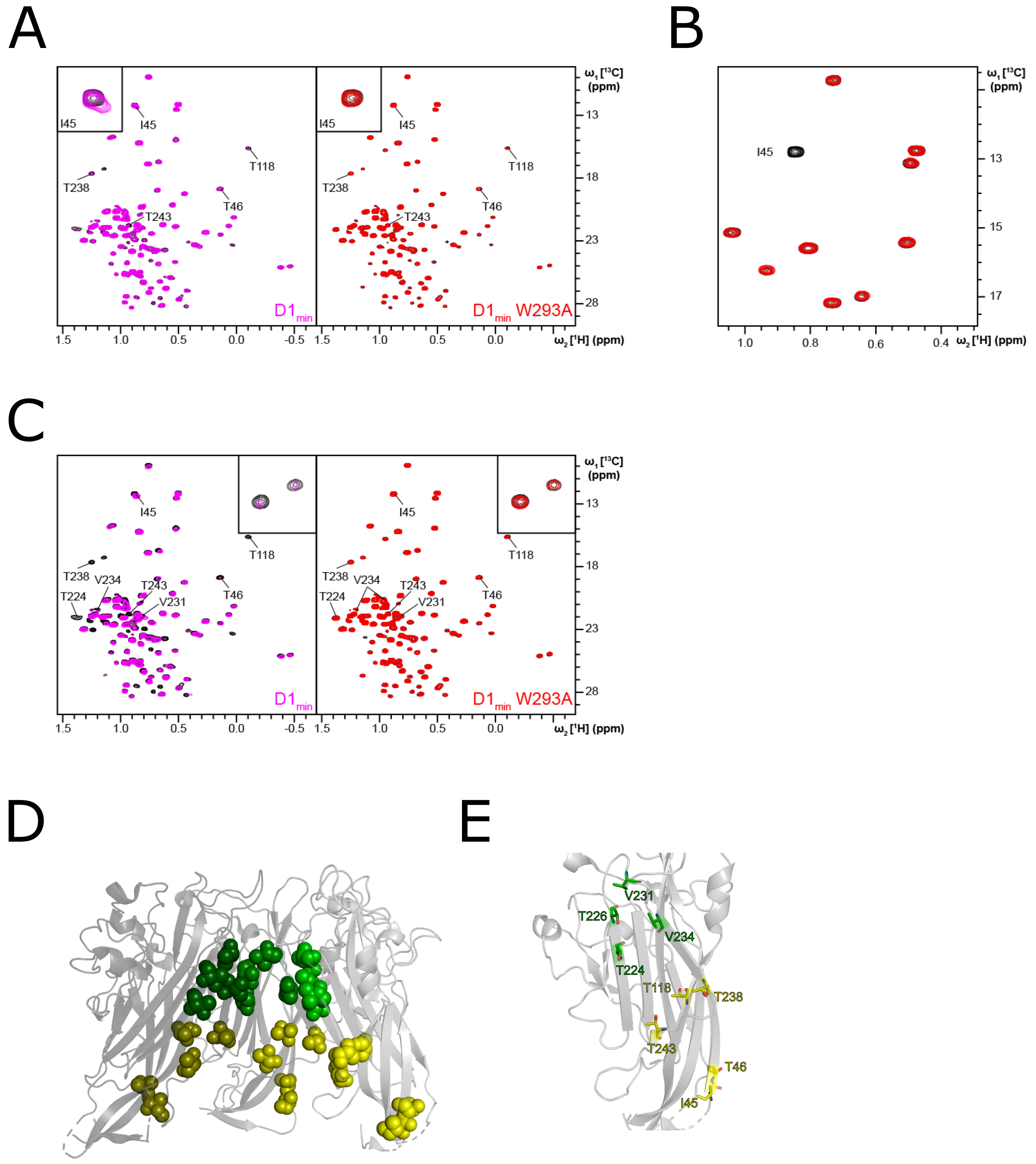

### Supplemental Figure S3

#### Supplemental Figure S3

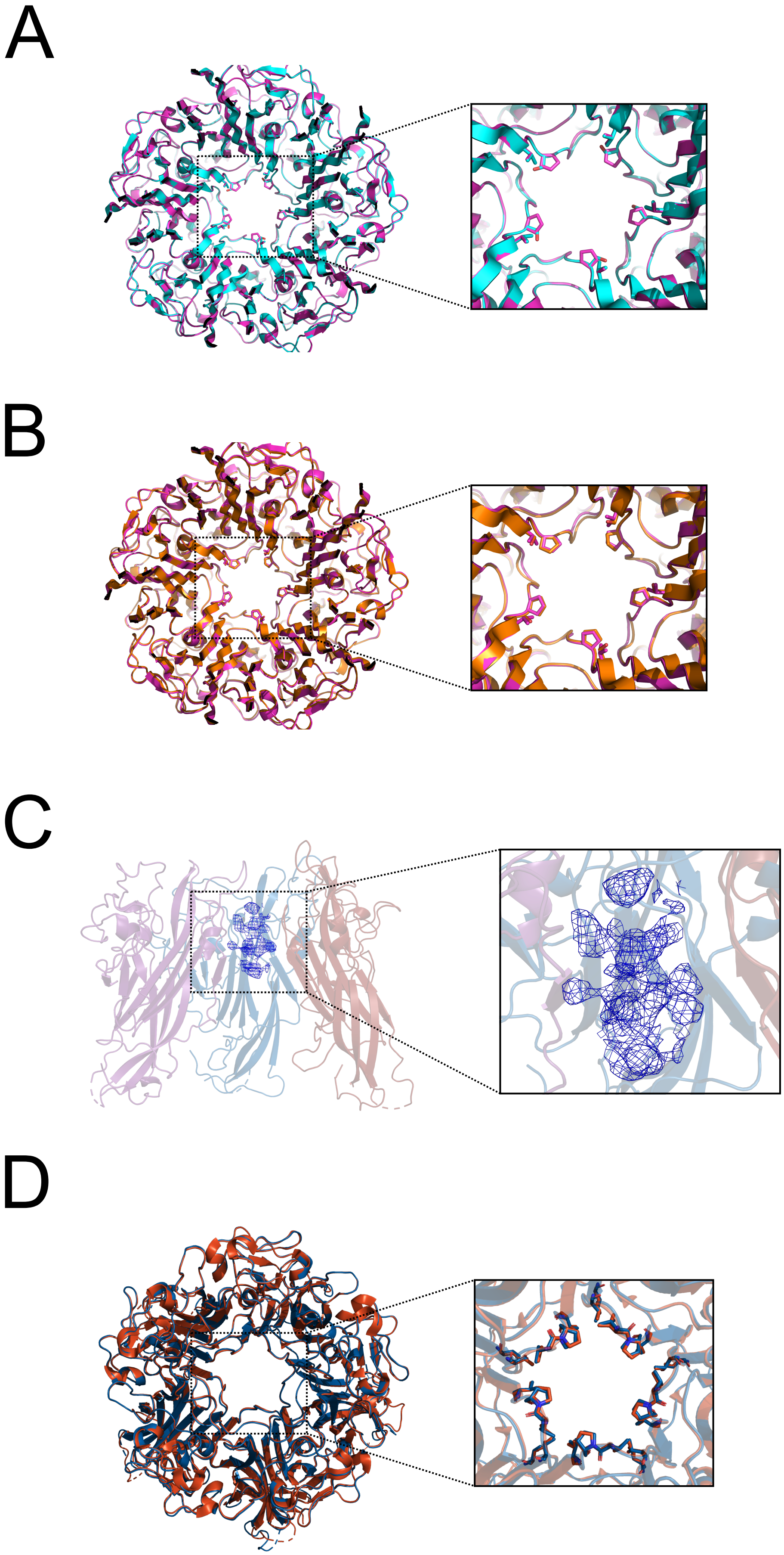

### Supplemental Figure S4

#### Supplemental Figure S4

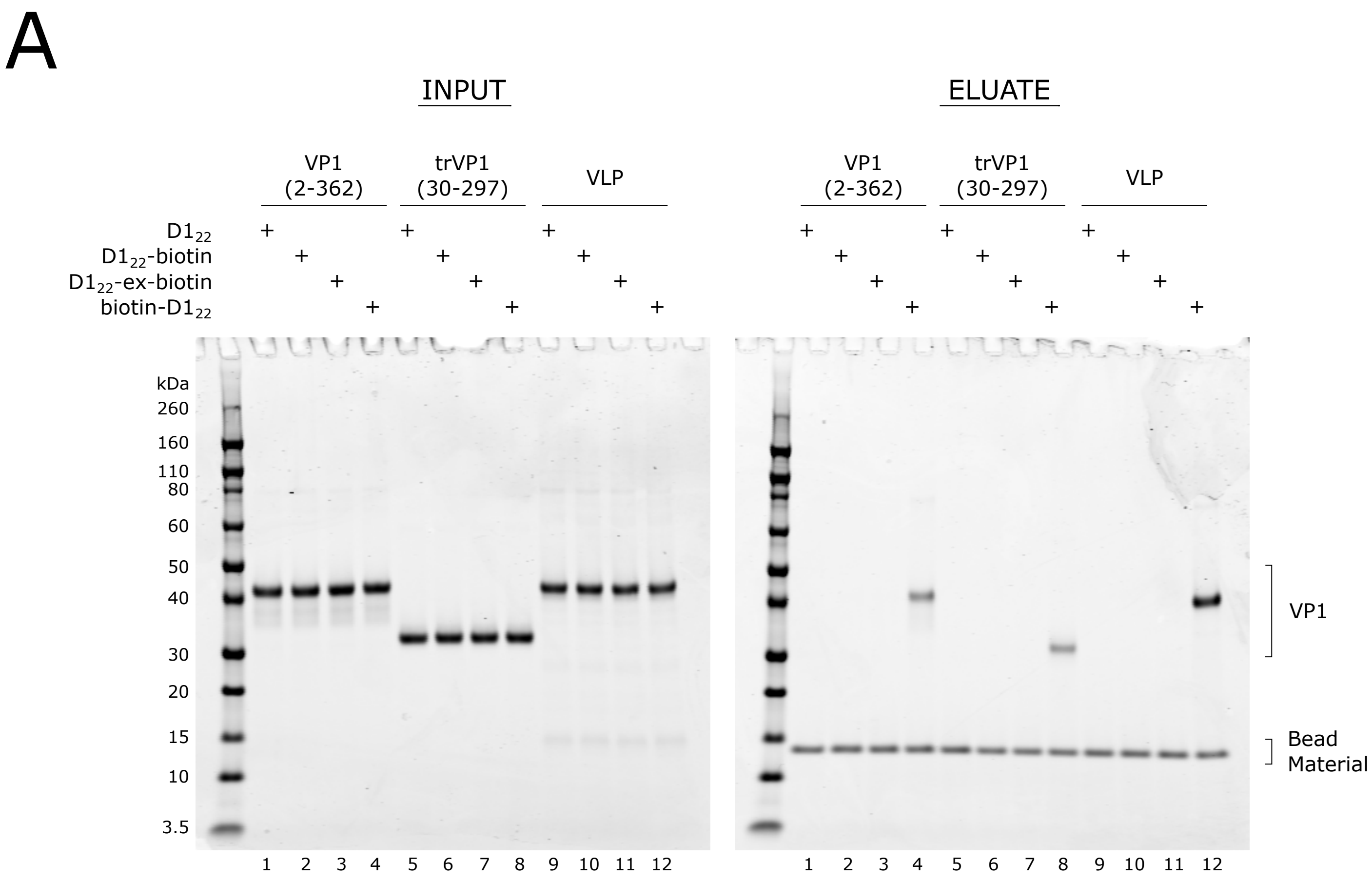

### Supplemental Figure S5

#### Supplemental Figure S5

A

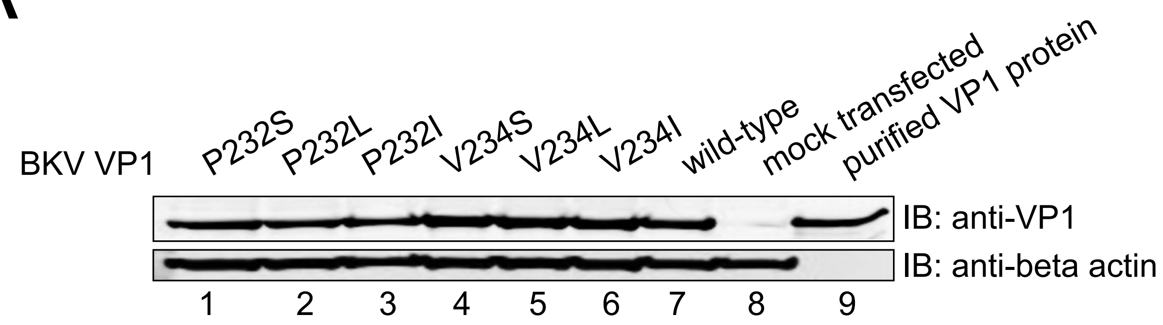


B

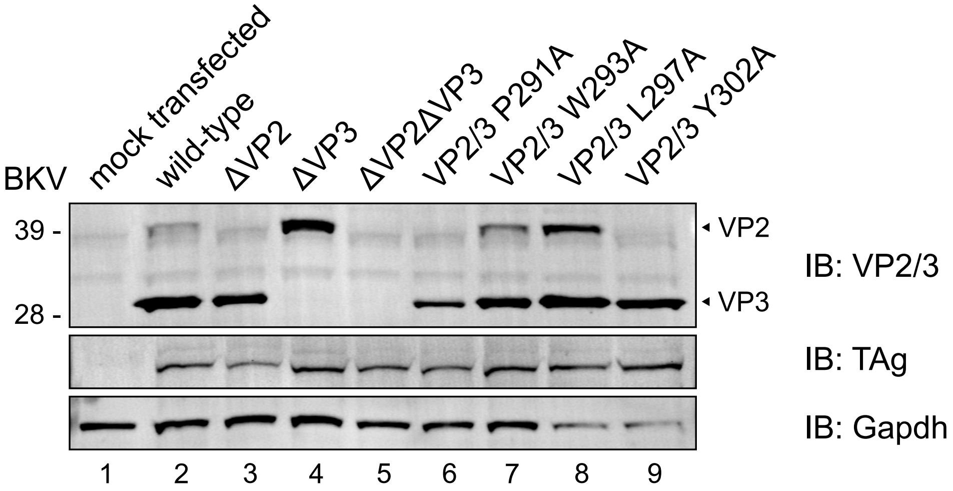
